# Modeling Parent-Specific Genetic Nurture in Families with Missing Parental Genotypes: Application to Birthweight and BMI

**DOI:** 10.1101/2020.08.06.239178

**Authors:** Justin D. Tubbs, Liang-Dar Hwang, Justin Luong, David M. Evans, Pak C. Sham

**Author notes:** Joint First Authors (Justin D. Tubbs, Liang-Dar Hwang). Joint Senior Authors (David M. Evans, Pak C. Sham). Correspondence: Pak C. Sham 1/F, Jockey Club Building for Interdisciplinary Research 5 Sassoon Road, Pokfulam, Hong Kong.

## Abstract

Disaggregation and estimation of genetic effects from offspring and parents has long been of interest to statistical geneticists. Recently, technical and methodological advances have made the genome-wide and loci-specific estimation of direct offspring and parental genetic nurture effects more possible. However, unbiased estimation using these methods requires datasets where both parents and at least one child have been genotyped, which are relatively scarce. Our group has recently developed a method and accompanying software (IMPISH; Hwang et al., 2020) which is able to impute missing parental genotypes from observed data on sibships and estimate their effects on an offspring phenotype conditional on the effects of genetic transmission. However, this method is unable to disentangle maternal and paternal effects, which may differ in magnitude and direction. Here, we introduce an extension to the original IMPISH routine which takes advantage of all available nuclear families to impute parent-specific missing genotypes and obtain asymptotically unbiased estimates of genetic effects on offspring phenotypes. We apply this this method to data from related individuals in the UK Biobank, showing concordance with previous estimates of maternal genetic effects on offspring birthweight. We also conduct the first GWAS jointly estimating offspring-, maternal-, and paternal-specific genetic effects on body mass index.

## Introduction

Renewed interest has been drawn towards estimating the effects of genetic nurture on offspring phenotypes using genome-wide single nucleotide polymorphism (SNP) data. Genetic nurture refers to the effects of parental genotypes on the phenotype of their child through influences on the child’s physical or social environment. Although geneticists have long criticized social scientists for inadequate control for the effects of genetic transmission when evaluating cultural transmission and the effects of parental behavior on children, geneticists themselves also often overlook the effects of cultural transmission (through genetic nurture) when evaluating genetic transmission. Specifically, genome-wide association study (GWAS) effect size estimates may be biased because of failure to consider parental effects on environmental factors relevant to the phenotype (Warrington et al. 2018; Kong et al. 2018).

Previous studies have attempted to quantify genetic nurture by calculating polygenic scores (PGS) of transmitted and un-transmitted alleles of each parent in parent-offspring trios, using effect size estimates from GWAS, and demonstrated genetic nurture effects of both maternal and paternal genotypes on offspring education attainment (Bates et al. 2018; Kong et al. 2018). However, these models are complicated by the need to determine the transmitted and un-transmitted alleles of each parent, which is not possible for some SNPs even in complete parent-offspring trios, and even more difficult when genotype data is missing for one of the parents. We recently showed that a conditional model which considers entire individual-specific PGS from offspring and their parents without partitioning into transmitted and non-transmitted components results in modest power gain in detecting genetic nurture, and used this approach to demonstrate a small role for maternal genetic nurture on offspring BMI trajectories (Tubbs et al. 2020a). By using traditional summary statistics to calculate polygenic scores, such models also implicitly assume that a variant’s genetic nurture effect has the same direction, and same relative magnitude, as its effect through genetic transmission.

Other groups have developed and applied methods to estimate the effects of child and maternal genotypes on offspring birthweight for specific loci through structural equation modeling which only requires genotypic information on the mother and phenotypic information on both the mother and one of her children (Warrington et al. 2018, 2019; Moen et al. 2019). These studies have provided important insight into the biological mechanisms involved in determining birthweight. For example, demonstrating opposing genetic nurture vs genetic transmission effects of high glucose risk alleles on offspring birthweight.

As studies on birthweight have shown, parental and offspring genetic effects can be discordant. Additionally, traditional GWAS estimates can be biased when parental genetic effects exist. Thus, it would be preferable to obtain conditional estimates of maternal, paternal, and offspring genetic effects to perform PGS-based methods as described previously. However, few GWAS have genotypic data on mothers and fathers, in addition to genotype data on and the index subject (the child), making it difficult to estimate or control for paternal and maternal effects, as well as the effects of the child’s genome. When genotype data of one parent is missing but has a true effect on the offspring phenotype, estimation of the genetic effects of both the remaining parent and the child are biased (Tubbs et al. 2020b). Since many family datasets have no genotype data on the father, such as the Raine Study cohort (Straker et al. 2017) or the Born in Bradford cohort (Wright et al. 2013), this raises the problem of bias when estimating maternal effects, except when paternal effects can be confidently assumed to be absent, such as for neonatal traits likely to be influenced by maternal intra-uterine environment. Motivated by this scenario, and noting that the UK Biobank (UKB; Bycroft et al., 2018) include a sizeable number of sibling relationships, we recently developed a method to detect maternal genetic effects on offspring phenotypes by imputing missing maternal genotypes from the genotypes on available full- and half-sibships (Hwang et al. 2020). We showed analytically and by simulation that association analysis of imputed maternal genotype gives unbiased estimates for effects of the mother’s and the child’s genotypes, provided that paternal effects are absent.

Since the assumption that paternal effects are absent is only reasonable for neonatal phenotypes, a method that allows discrimination of maternal and paternal effects is highly desirable. We noted that mother-child and father-child pairs are often more frequent than complete parents-child trios in GWAS, and hypothesized that such pairs would allow unbiased estimation of the effects of the mother’s genotype, the father’s genotype, as well as the effects of the child’s genotype. Here, we show that association analysis using the imputed genotype of the missing parent, through closed-form derivation and simulation, results in asymptotically unbiased estimates of maternal, paternal, and offspring genetic effects. We derive the statistical power characteristics of these models and implement them in a convenient online power calculator (https://evansgroup.di.uq.edu.au/power-calculators.html). Application of this method to family data from the UKB shows concordance with previous estimates of maternal genetic effects on offspring birthweight. Finally, we applied the method on a genome-wide scale to perform the first GWAS of maternal and paternal genetic effects on offspring body-mass index (BMI).

## Methods

### Overview

The proposed method jointly estimates the regression coefficients of a child’s quantitative phenotype on the genotypes of the child, the father and the mother, from nuclear family data, where the genotype is coded additively as 0, 1 and 2 – the allele dosage for an autosomal bi-allelic locus. When some nuclear families contain two or more children with measured phenotypes, a random effect is introduced to take account of residual phenotypic correlations between children of the same family. For nuclear families where the genotype of one parent is missing, the missing parent’s genotype is imputed by the conditional expectation of the genotype, given the available genotypes of family members. This conditional expectation is calculated from the conditional probabilities of the possible genotypes of the missing genotype by Bayes’ theorem. These families are then analyzed together with families where genotypes are available for both parents under a linear regression model, or a mixed model if some families have two or more children with measured phenotypes.

Singletons where neither parental genotype is observed are excluded from analysis, as the conditional genotype of a parent would be collinear with the genotype of the child. When neither parental genotype is observed, but the genotypes of two or more children are available, the conditional expectations of the two parents, while not collinear with the genotypes of either child, are identical to each other. Thus, we can only impute a “parental” genotype, and the regression coefficient of this imputed parental genotype would represent the sum of the effects of the two parents. To avoid the need to impose constraints on parameter estimates, we analyze these families separately from those with one or no missing parents. The summary statistics from the analyses of these two distinct datasets are then combined to provide overall estimates of child, paternal and maternal effects.

### Statistical model

Our model assumes random mating and the absence of dominance. The offspring, maternal and paternal genotypes *X*_*o*_, *X*_*m*_ and *X*_*f*_ respectively, influence an offspring phenotype *Y*_*o*_ through both genetic and cultural transmission.

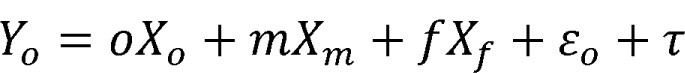

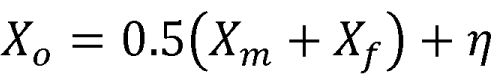

where *o* is the effect of the offspring genotype (*X*_*o*_), *m* is that of the mother’s genotype, and *f* is that of the father’s genotype. The random variable *ε*_*o*_ accounts for residual phenotypic variance, and *η* accounts for half the variance of the offspring genotype – the segregation variance. The random variable *τ* captures the random effect for family when there is more than one offspring observed, accounting for the residual correlations between siblings.

### Imputation procedure and precision

To illustrate why parental imputation is possible, consider a biallelic locus where the observed offspring has genotype *Aa* and the observed mother has genotype *AA*. The father must then have genotype *Aa* or *aa*, and under random mating, these two possible genotypes have probabilities that depend only on the relative frequencies of alleles *A* and *a* in the population. If another offspring of these two parents is observed to have *AA*, then we can impute the father’s genotype with certainty as *Aa*.

The imputation of a parental genotype in nuclear families with 2 or more offspring has been described in detail elsewhere (Hwang et al. 2020). The method imputes a missing parental genotype as the conditional mean of the possible genotypes of the parent, where the probabilities of the possible genotypes are calculated by Bayes’ theorem. When applied to impute a missing maternal genotype from the observed genotypes of the father and an offspring, the Bayes’ formula can be written as follows for a nuclear family:

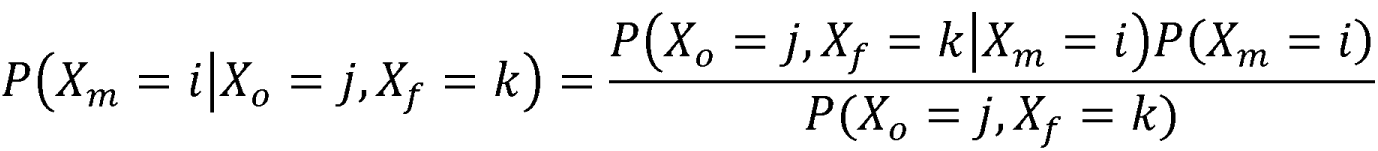

where *i* contains the set of possible genotypes for the (unobserved) mother, while *j* and *k* are the observed genotypes of the offspring and father, respectively.

Imputed parental genotypes are generally less variable than the true underlying parental genotypes. The accuracy of an imputed parental genotype, say 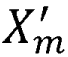, can be measured by the variance of 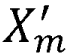 relative to the variance of *X*_*m*_. We have derived imputation accuracy for different family configurations, as a function of allele frequency. For example, the imputation accuracy of a maternal genotype from the observed genotypes of the father and an offspring, for a locus with allele heterozygosity *H* (=2*p*(1-*p*), where *p* is allele frequency), is (2*H* – *H*^2^)/4. The imputation variances and accuracy for different family configurations and allele frequencies are plotted in Figure 1.

**Figure 1.**
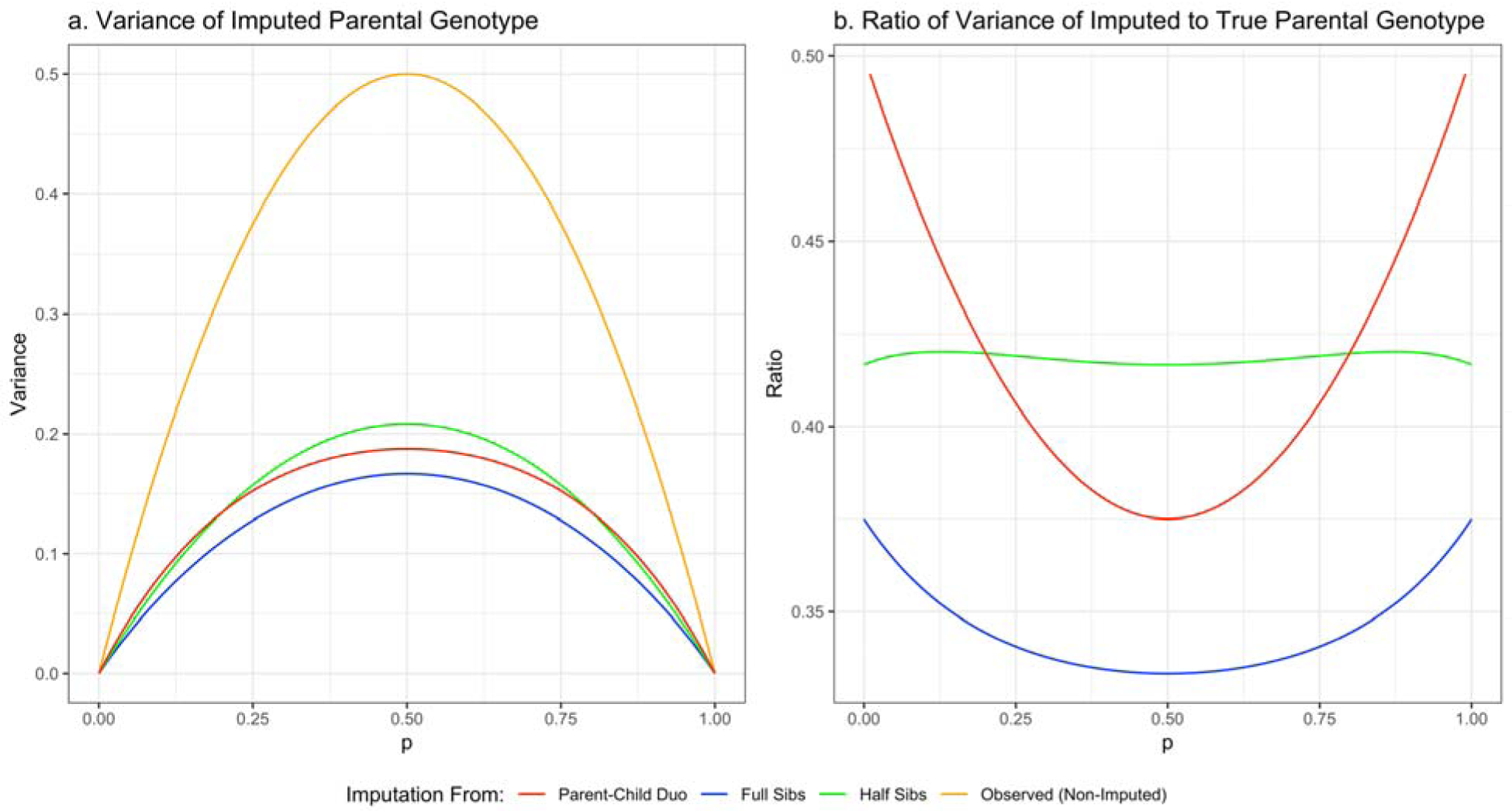
Variance of Imputed Parental Genotype. *Panel a shows the expected variance of an observed parental genotype for various values of minor allele frequency, p, compared the variance of the parental genotype imputed from various relationship types. Similarly, panel b compares the imputation accuracy, measured as the ratio of imputed to true (observed) variance, across different imputation sources. Note that half sibs under this model share the parent which is imputed*.

Since the imputed parental genotype 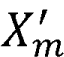 is a conditional expectation of the underlying parental genotype *X*_*m*_, we can infer that the regression coefficient of *X*_*m*_ on 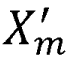 is 1. It follows that 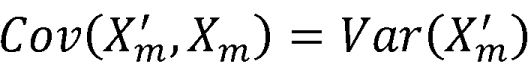. If the regression coefficient of phenotype *Y* on *X*_*m*_ is *β*, then the regression coefficient of *Y* on 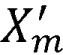 is 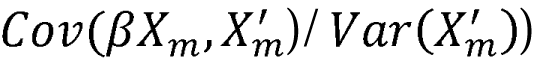, i.e. also *β*. This provides an intuitive explanation of why the regression of the phenotype on an imputed parental genotype provides an unbiased estimate of the regression of the phenotype on the underlying parental genotype, which can also be visualized as a path diagram in Figure 2.

**Figure 2.**
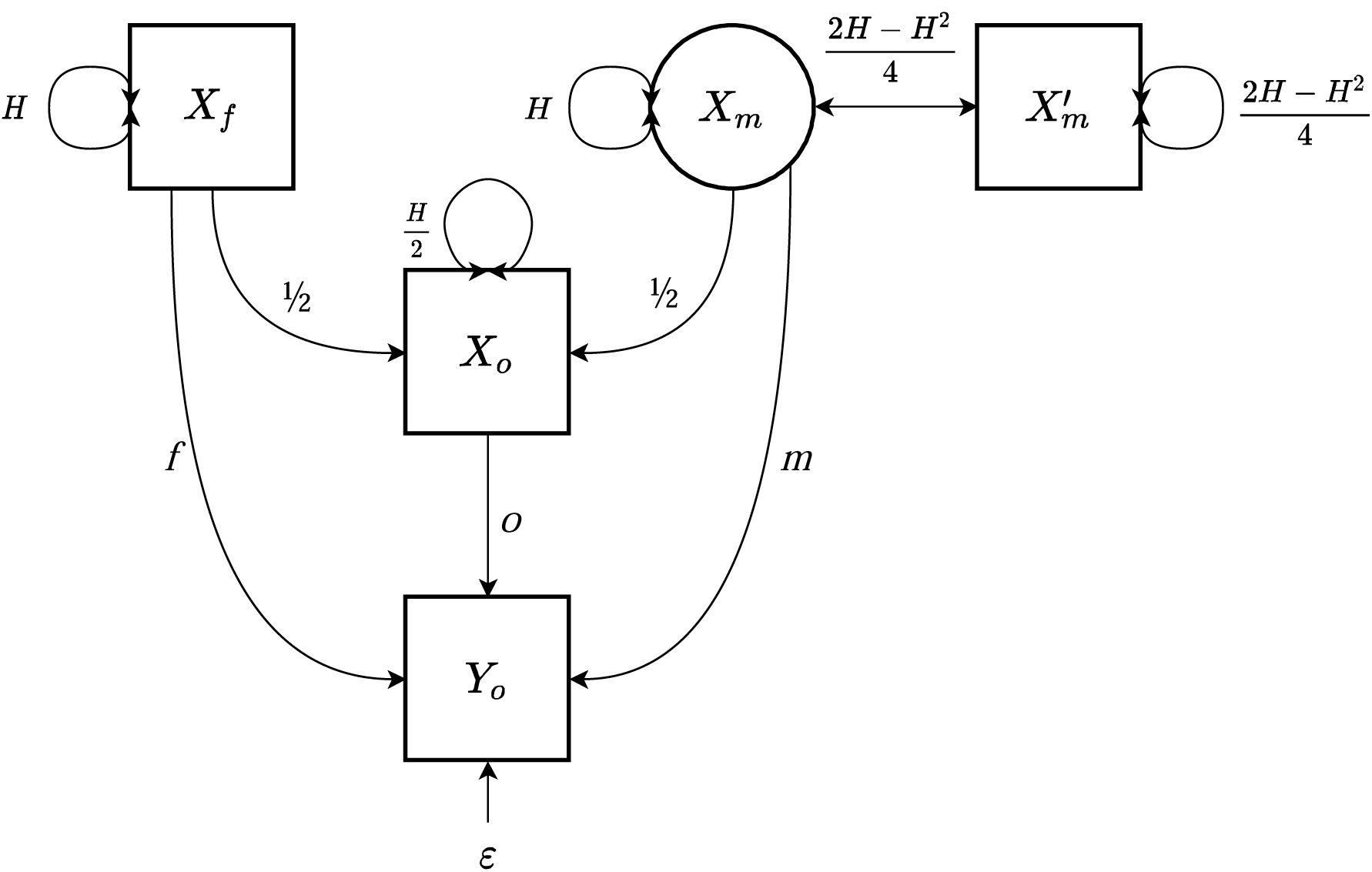
Path Diagram illustrating estimating the unbiased effect of maternal genotype on offspring phenotype using imputed maternal genotype. *This path diagram illustrates the situation where the offspring phenotype, Y_o_, and genotype, X_o_, are observed along with the paternal genotype, X_f_. The mother’s genotype is imputed as described in the text as X′_m_. X_m_ represent the unmeasured true maternal genotype. The child’s genotype has direct effect o, the paternal genotype has effect f, and the maternal genotypic effect is represented by m. ε is the residual random error variance in the offspring phenotype. H is the heterozygosity, 2p(1-p), where p is the minor allele frequency*.

### Overall parental effect estimates

Two separate mixed model analyses are performed: (1) for families with 0 or 1 missing parent, (2) for families with 2 missing parents. From (1) we obtain separate estimates for both paternal and maternal effects, whereas from (2) we obtain only an estimate for the sum of the paternal and maternal effects. Noting that separate paternal and maternal effects can be transformed to a sum and a difference, and vice versa, we first first transformed the separate paternal and maternal effect estimates from (1) into a sum and a difference, so that the sum can be combined with the total parental effect estimate from (2). The resulting overall total parental effect estimate, together with the difference, can then be transformed to separate overall paternal and maternal effect estimates. The detailed procedure is described in the Supplementary Material.

### Analytic power calculation

In the Supplementary Material, we present this model in further detail and provide a complete derivation of the asymptotic non-centrality parameter (NCP) and bias characteristics of a multiple linear regression model where the offspring phenotype is regressed on the child’s, mother’s, and father’s genotypes. Furthermore, we consider the situation where the father’s genotype is missing and is imputed as its conditional expectation given the observed mother-offspring duo and the population allele frequency. Finally, we detail our tailor-made procedure (described fully in the Supplementary Material) based on inverse covariance weighting to combine the estimates from models using imputed parental genotypes when 0, 1, or 2 parents are missing.

### Simulations

To confirm the accuracy of our derivations and combination method, we performed two sets of simulations using using R version 3.6.1 (R Core Team 2019). First, to ensure our power and bias calculations were accurate, we simulated a dataset of 10,000 trio family units genotyped at a single locus with a minor allele frequency of 0.25. Imputed paternal genotypes were calculated based on the offspring and maternal values following our method. We repeated this procedure, simulating offspring phenotypes for five different child-maternal-paternal effect size combinations using their observed genotypes, with 10,000 replicates for each effect size combination. Finally, we performed two regression analyses for each effect size combination – one where all genotypes are “observed” and included as predictors of the offspring phenotype, and one where the paternal genotype is replaced by its imputed genotype. We obtained the regression estimates for each genotypic effect and calculated power and type I error rate, comparing these to the results from our derived formulae.

A second simulation procedure was performed to show that our combination method produces unbiased estimates of maternal, paternal, and offspring genetic effects in a sample with similar characteristics to the UKB. We simulated a dataset which approximates the structure of available family data in the UKB with 20,000 sibling-pairs and 5,000 singletons along with the genotypes of their parents. Similar to the previous simulation, we varied the size of the offspring, maternal, and paternal genetic effects across five combinations, while maintaining a minor allele frequency of 0.25 and a residual phenotypic correlation of 0.2 between siblings. For each effect size combination scenario, we performed 10,000 simulation replicates applying our proposed analytic strategy by constructing one linear model for each data type.

For trios, where a child and both parents are genotyped, we performed a multiple linear regression to jointly estimate the effects of offspring, maternal, and paternal genotypes on the offspring phenotype. For sibling pairs, we used the lme4 package in R (Bates et al. 2015) to estimate a linear mixed model with a random effects term to account for within sibling variance and fixed effects terms for the offspring genotypic effect and that of the combined parental genotype (sum of mother and father). From the output of these models, we applied our proposed inverse covariance weighting strategy to obtain combined estimates of offspring, maternal, and paternal genotypic effects. These models treated all parental genotypes as if they had been observed, which would be the best-case scenario.

To approximate the pattern of parental genotype missingness in the UKB, we treated all sibling parental genotypes as if they were unobserved and used the method from Hwang et al. (2020) to impute the parental genotype for these sibling pairs. Similarly, for trios, we treated 30% of the sample as if the mother had been unobserved and 60% as if the father had been unobserved, imputing their genotypic value following our proposed method. Again, we applied our proposed linear modeling procedure to these data separately for trio data (with one or no missing parents) and sibling data (with both missing parents) and combined their estimates using the method described in the Supplementary Material. We then compared the distribution of parameter estimates across 10,000 replications from the linear models where all parents are observed versus the second situation where the parental genotype availability approximates that of the UKB and imputed using our proposed method.

### Application

We applied our imputation model genome-wide to birthweight and BMI for nuclear families in the UKB. First, participants of white British ancestry who had at least one family member also in the UKB, whose genetic sex matched their reported sex, and who passed the genotyping quality control filters set by UKB (Bycroft et al. 2018) were identified. We then applied the relationship inference criteria recommended by the authors of KING (Manichaikul et al. 2010) along with information on year of birth (YOB) and sex to the output from this software provided by UKB to classify first-degree relative pairs as siblings or parent-offspring, excluding monozygotic twins. These pairs were then clustered into nuclear families to identify complete families, parent-child duos, and sibships using the igraph R package (Csardi and Nepusz 2006). For each nuclear family at each genotyped SNP with a minor allele frequency greater than 0.001, the missing parent(s) were imputed following the applicable method as described in Hwang et al (2020) or in the Supplementary Material using the genotype of all available first-degree relatives and the effect allele frequency as calculated by PLINK version 1.9 (Chang et al. 2015) from the full sample of white British participants in the UKB passing their genotyping quality control filters (Bycroft et al. 2018).

For those who had phenotypic values from more than one clinical assessment, the average of all available reports was taken. Offspring were excluded from analysis of birthweight if any two self-reported own birthweight values differed by more than 0.5kg or if their reported birthweight was less than 2.5kg or greater than 4.5kg, which are unlikely values for this cohort (Warrington et al. 2018). The mean birthweight for offspring in our analyzed sample was 3.39kg (SD = 0.42). BMI was transformed by taking the natural logarithm of each raw BMI value plus 3, which was found to minimize the skew of the resultant distribution. The average offspring BMI for our sample prior to transformation was 27.22 (SD = 4.74). The available sample sizes for the imputation procedure and phenotypic analyses by family type are shown in Table 1.

**Table 1.**
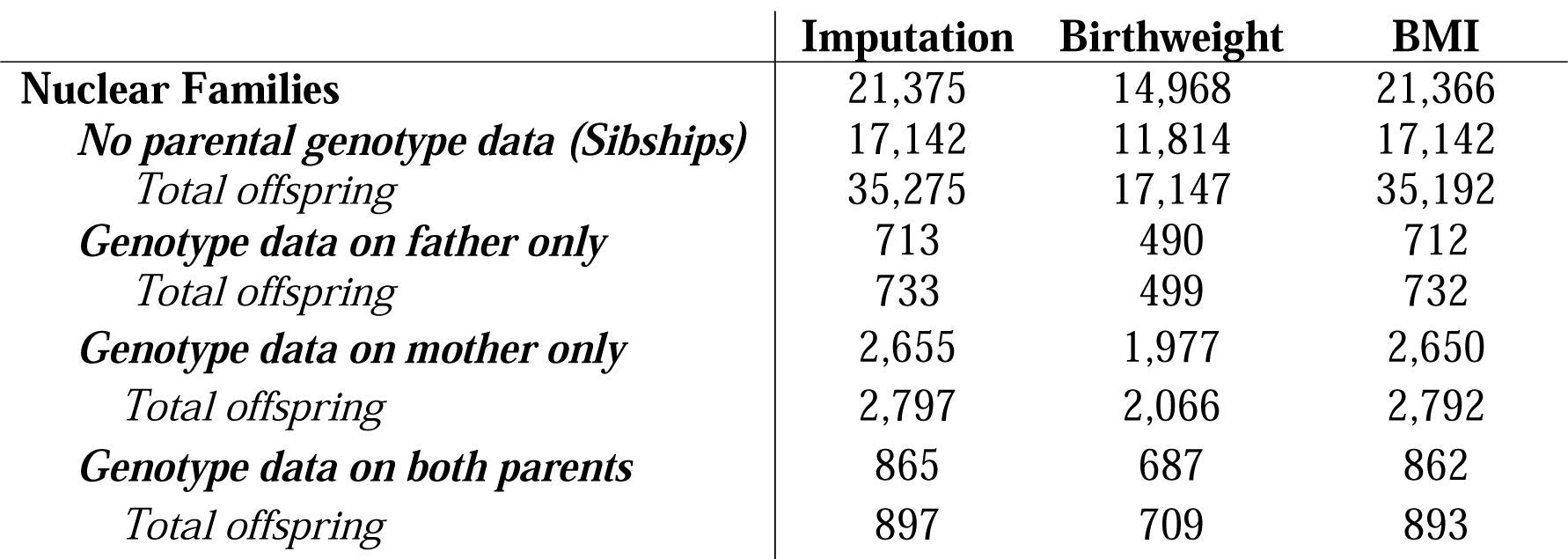
Sample Size by Family Type for UK Biobank Analyses. *This table shows the sample characteristics for our imputation procedure, as well as for analyses of birthweight and BMI in the UK Biobank. These values reflect the maximum analytic sample size for any SNP after filtering the sample as described in the text*.

|  | **Imputation** | **Birthweight** | **BMI** |
| --- | --- | --- | --- |
| **Nuclear Families** | 21,375 | 14,968 | 21,366 |
| *No parental genotype data (Sibships)* | 17,142 | 11,814 | 17,142 |
| *Total offspring* | 35,275 | 17,147 | 35,192 |
| *Genotype data on father only* | 713 | 490 | 712 |
| *Total offspring* | 733 | 499 | 732 |
| *Genotype data on mother only* | 2,655 | 1,977 | 2,650 |
| *Total offspring* | 2,797 | 2,066 | 2,792 |
| *Genotype data on both parents* | 865 | 687 | 862 |
| *Total offspring* | 897 | 709 | 893 |

For each phenotype, two mixed models were run at each genotyped SNP using the lme4 package (Bates et al. 2015) in R with a random effect term included to account for residual covariance within families when two or more siblings were observed. The outcome and all non-genetic predictor variables were standardized to optimize the model estimation procedure and running time of lme4. The first mixed model was applied to data on trios (i.e. 0 or 1 missing parent) and included fixed effects terms for the additively coded child SNP, maternal SNP, paternal SNP, YOB, YOB^2, sex, assessment center, and the first 10 genetic principal components as provided by the UKB. The maternal and paternal SNPs were included if observed, or substituted with their imputed values if originally missing. The second mixed model was applied to data on siblings (i.e. both parents missing) and was equivalent to the first model except that instead of separate maternal and paternal genotype predictors, the imputed genotype, which represents the combined effects of both parental genotypes was included. Subsequently, the estimates from these two models were combined using the method described in the Supplementary Material. P-values were subsequently calculated for these combined estimates. For the model predicting birthweight, the paternal genotype was not included as a predictor. It is less likely that the father’s genotype would have a large effect, if any, on the offspring’s birthweight other than through genetic transmission. Since we expect this effect for most SNPs to be zero, by not estimating the father’s genotypic effect, we effectively increase power to detect other genetic effects while still obtaining asymptotically unbiased estimates (Tubbs et al. 2020b).

Although our analysis is the first of its kind for BMI, previous studies have estimated the conditional effects of offspring and maternal genotypes on birthweight, indicating a strong influence of metabolic traits mediated through the maternal genotype (Beaumont et al. 2018; Warrington et al. 2018, 2019). Thus we compare the genetic effect estimates from our model with those reported by Warrington and colleagues (2019) for 155 SNPs which were observed in our sample or for which suitable proxy variants (R^2^>0.9) could be found using the SNiPA proxy search tool (Arnold et al. 2015). Additionally, given the link between diabetes, maternal BMI, and offspring birthweight (Ornoy 2011; Tyrrell et al. 2013; Kong et al. 2019), we performed targeted analyses using genome-wide significant loci identified in recent GWAS of type-II diabetes (Xue et al. 2018) and BMI (Yengo et al. 2018) as candidate SNPs to test for maternal genetic nurture effects at these loci. For each of the two GWAS summary statistics, we identified genome-wide significant SNPs which overlapped with those in our sample. We then applied a Bonferroni-corrected p-value threshold based on the number of independent loci identified in the respective study to identify loci with a potential maternal genetic effect on birthweight in our sample (i.e. 0.05/143 for diabetes and 0.05/941 for BMI). Finally, we performed the same candidate SNP testing procedure to the results from our conditional analysis of BMI, testing for maternal and paternal genetic effects of overlapping SNPs identified as significant in the GWAS by Yengo et al. (2018) using a Bonferroni-corrected p-value threshold of (0.05/941).

## Results

### Analytic Derivation and Simulations

The full derivation of beta estimates and NCPs under a linear regression model for observed and imputed genotypes are detailed in the Supplementary Material. We show that unbiased estimates of offspring and parental genotypic effects are obtained even when one parent (e.g. the father) is ungenotyped and imputed using our proposed method as

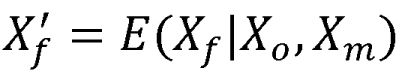

Table 2 shows the asymptotic power characteristics of our imputation regression model compared to the situation where either all genotypes are observed, or the missing parent is simply unmodeled (as derived in Tubbs, Zhang, & Sham, 2020). A convenient online power calculator is available at https://evansgroup.di.uq.edu.au/power-calculators.html. Although our imputation model addresses the bias which would be introduced by excluding the missing parental genotype, it also results in decreased power as compared to the most ideal situation where all genotypes are observed.

**Table 2.**
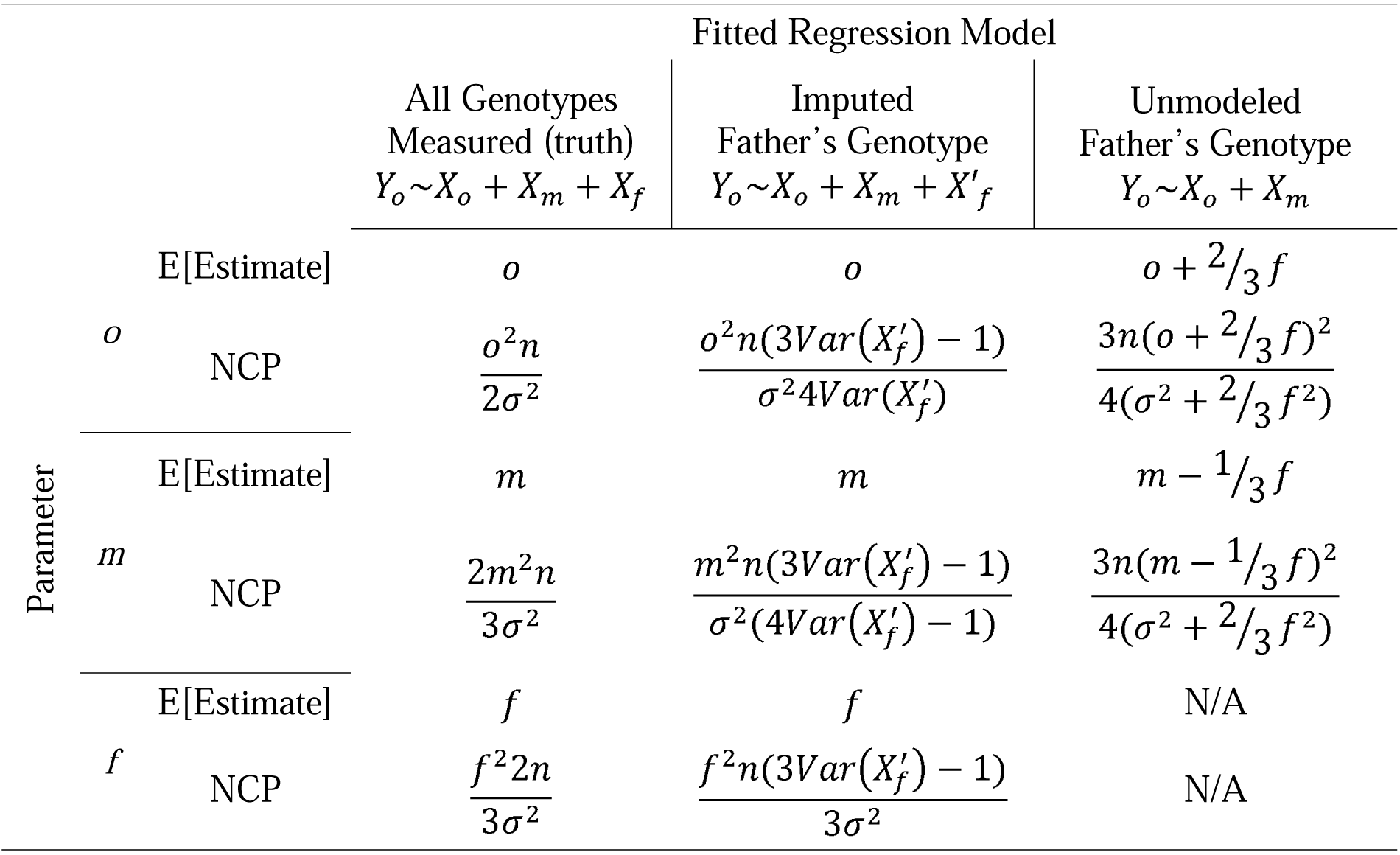
Bias and NCP of Regression Estimates. *This table presents derived regression estimates and non-centrality parameters for linear regression models predicting the child’s phenotype (*Y*_*o*_) across three situations: when both parents’ genotypes are observed and included in the model, when one parent is missing and its imputed value is included in the model, and when one parent is missing and not included in the model (from Tubbs, Zhang, & Sham, 2020). Unbiased estimates can be recovered through our imputation procedure, albeit at the expense of statistical power. o is the direct effect of the offspring’s genotype on the offspring’s phenotype, m is that of the mother’s genotype, and f is that of the father’s. n is sample size and σ^2^ is the error variance in the child’s phenotype under the given mode. Y_o_ is the child’s phenotype, X_o_ is the child’s genotype, X_m_ is the mother’s genotype, X_f_ is the father’s, and X′_F_ represents the father’s imputed genotype. E[Estimate] is the expectation of the regression beta estimate*.

Supplementary Figure 1 shows the derived statistical power to detect an offspring, maternal, or paternal genetic effect at a single locus under a one degree of freedom chi-square test at an alpha level of 0.05 across different values of true paternal effect when all three genotypes are observed versus when all paternal genotypes are imputed from genotyped mother-offspring duos. Interestingly, the power to detect a maternal or offspring genetic effect is independent of the true value of the paternal effect. However, when the father’s genotype is imputed, the power to detect a maternal of offspring effect is decreased as a function of the imputed paternal variance (which itself is a function of the minor allele frequency).

Simulation results are provided in Supplementary Tables 1 and 2. Simulation 1 confirms the accuracy of our analytic power derivations, with empirical power similar to the predicted values for all effect size combinations. Regression beta estimates for offspring, maternal, and paternal genetic effects are unbiased about their true simulated value, and type-1 error is adequately controlled at the specified alpha significance level, even when the father’s genotype is imputed. Similarly, simulation 2 shows that our combination method provides unbiased estimates in a sample with relationship types and missingness structure similar to that of the UKB. Figure 3 visualizes these results, showing that our model using imputed genotypes is able to provide unbiased estimates about the simulated true value.

**Figure 3.**
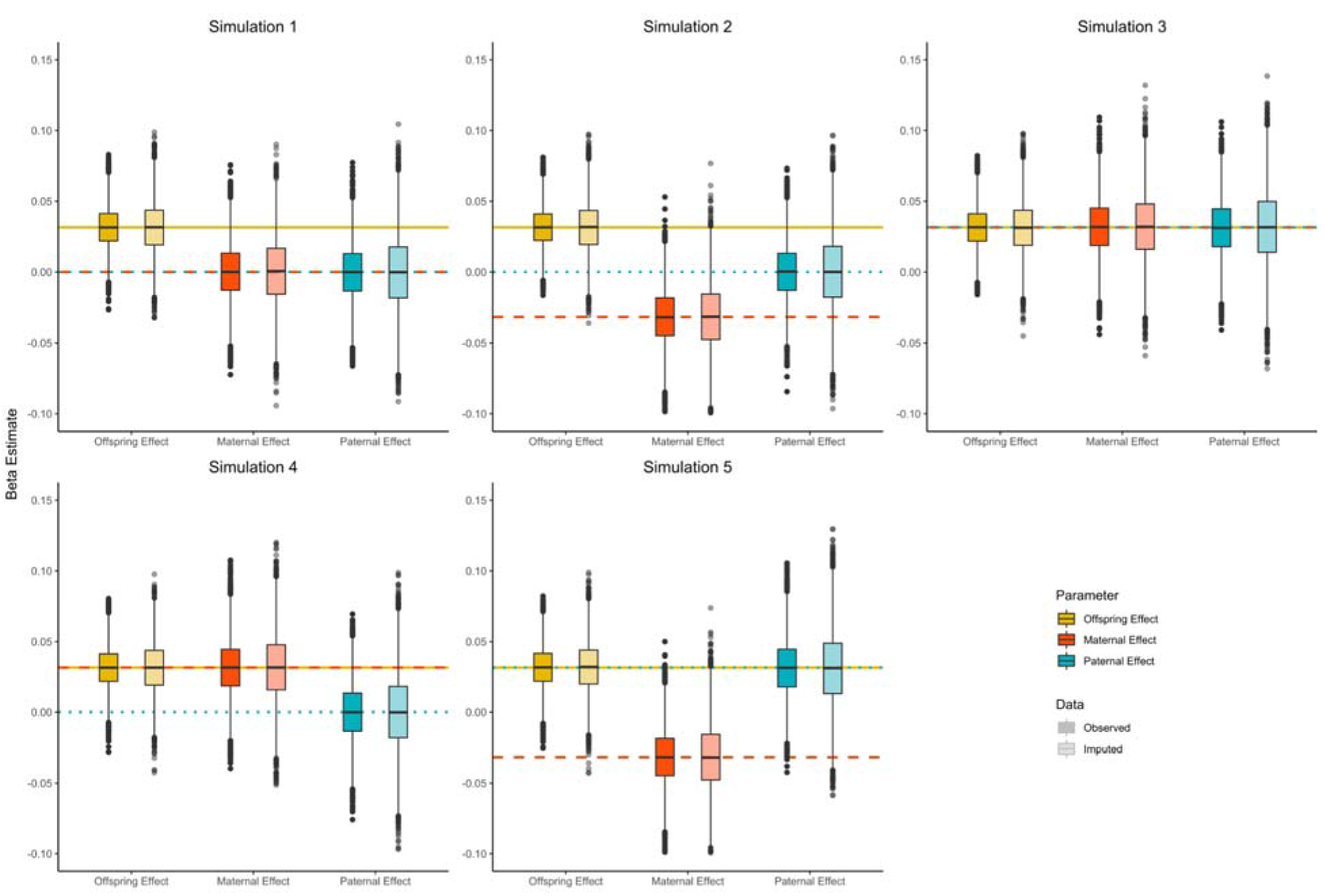
Simulation Beta Estimates. *Note: This figure shows unbiasedness of our method for estimating offspring, maternal, and paternal genetic effects for a single locus across five simulation parameter situations with 10,000 replicates each. The data are simulated to approximate the structure of available family data in the UK Biobank as detailed in the main text, while the three effect size combinations vary across each situation, explaining 0 or 0.1% of the variance in the offspring phenotype. The true effect sizes used to generate the phenotype are plotted as colored horizontal lines. The Observed Model is that where the phenotype is regressed on the simulated offspring, maternal, and paternal genotypes. The Imputed Model is that where the parental genotype has been imputed for all sibs, and a proportion (resembling those in the UK Biobank) of maternal and paternal genotypes have been imputed from corresponding parent-offspring duos*.

### Application

We applied our imputation and estimation procedure to offspring birthweight and BMI for nuclear families in the UKB. Summary statistics for birthweight and BMI at all genotyped SNPs passing quality control filters are provided in Supplementary Tables 3 and 4, respectively. No SNPs surpassed the p-value threshold for genome-wide significance for birthweight. Supplementary Figure 2 shows the Manhattan plots for offspring and maternal genetic effects on offspring birthweight, while Supplementary Figure 3 shows their QQ plots.

We compared the estimates of offspring and maternal genetic effects on birthweight obtained from our model which utilized both imputed and observed parental genotypes to those obtained from a recent analysis including unrelated UKB participants which utilized a structural equation modeling framework (Warrington et al. 2019). The conditional maternal and offspring SNP effect estimates obtained from Warrington and colleagues are for known birthweight-associated SNPs identified in GWAS of unrelated subjects. Both maternal and offspring effect estimates from our imputation model correlated with these previous estimates, as shown in Figure 4.

**Figure 4.**
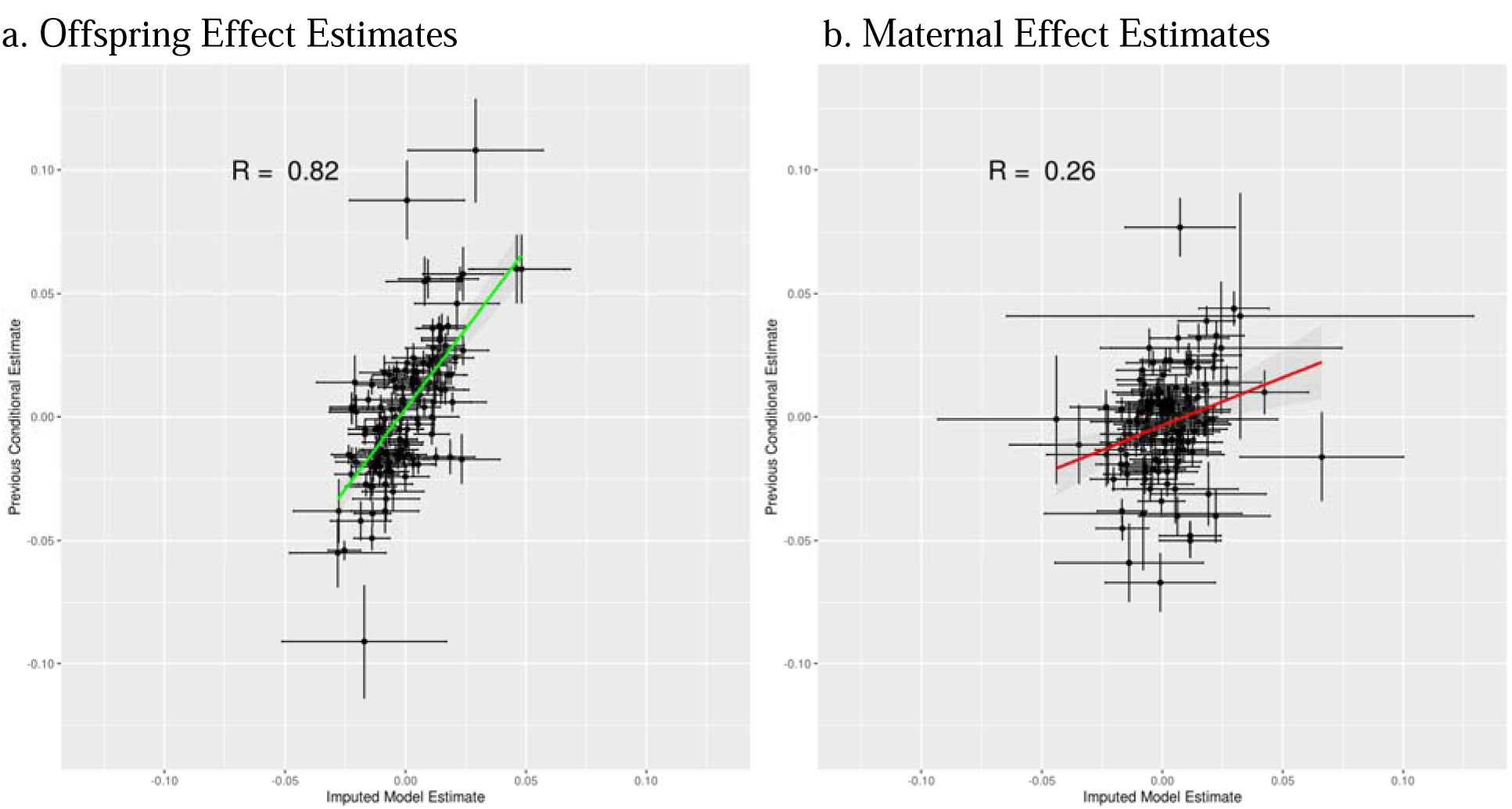
Offspring and Maternal Genetic Effects on Birthweight Estimated by our Imputation Model vs Previous SEM-Based Estimates in an Independent Sample. *We compare the estimates of offspring and maternal genetic effects on offspring birthweight obtained from our imputation model with those of a previous study employing a structural equation modeling framework (Warrington et al., 2019) across 155 SNPs which were observed in our sample or for which suitable proxy variants could be identified (R^2^>0.9). Warrington et al identified these as candidate SNPs, being identified in prior traditional GWAS as significantly associated with birthweight. Our conditional model, as described in the text, was applied to the UK Biobank, combining data from complete mother-child duos, father-child duos with imputed missing mothers, and sibships with imputed parental genotypes. As our model was performed on a scaled birthweight variable, we transformed our effect estimates and their standard errors to the original scale (for ease of comparison with the estimates from Warrington et al) by multiplying each estimate by the standard deviation of birthweight in our sample*.

As the power to detect parental effects at the traditional genome-wide significance level using these models is expected to be low, we also performed candidate SNP analyses based on significant SNPs identified through GWAS of type II diabetes and BMI to test for maternal genetic effects on offspring birthweight. These results are provided in Supplementary Tables 5 and 6. None of the candidate SNPs for diabetes passed the Bonferroni-corrected p-value threshold for the presence of a maternal genetic nurture effect. One candidate BMI SNP, rs3212335, passed the corrected p-value threshold for a maternal genetic effect on offspring birthweight with p = 7e-6. This SNP is an intronic variant in the *GABRB3* gene, which encodes a number of protein subunits of the GABA_A_ receptor. *GABRB3* is pivotal for proper neurodevelopment in humans, and shows parent-of-origin imprinting effects for certain disorders and in some cell types (Tanaka et al. 2010).

The GWAS results for maternal, paternal, and offspring genetic effects on BMI for families in the UKB are shown in Supplementary Figure 4, while Supplementary Figure 3 shows their QQ plots. SNPs within the well-known *FTO* gene indicated a genome-wide significant offspring genetic effect on BMI, but no genome-wide significant signals were observed with maternal or paternal genetic effects. One intergenic SNP on chromosome 9, rs17569929, approaches genome wide significance with a corrected p-value of 3e-8 for a maternal effect on offspring BMI, but this is likely to be a false positive given the absence of other significant variants around this loci. Likewise, no candidate SNPs for BMI indicated a maternal of paternal effect at the appropriate significance level, accounting for the number of independent BMI loci identified previously (0.05/941). These results are presented in Supplementary Table 7.

## Discussion

We have shown through analytic derivation and simulation that our imputation method can recover unbiased estimates of offspring, maternal, and paternal genetic effects using data from nuclear families, even when one or more parents have not been genotyped. This increases the utility of datasets where one or both parents have not been genotyped by increasing the effective sample size and allowing for parental effect estimation or control.

Furthermore, we demonstrate the utility of our method by applying the imputation and modeling procedure in a mixed model GWAS framework to sibships and parent-offspring families in the UKB. We show strong agreement between our method and previous estimates of offspring and maternal genetic effects on birthweight and through candidate SNP analysis, identify a locus in the *GABRB3* gene which may have a maternal genetic effect on offspring birthweight. We also perform the first GWAS to partition offspring and parental genetic effects on offspring BMI, but find no SNPs with significant maternal or paternal effects, even in a targeted analysis of candidate SNPs. Parental genetic effects on BMI are likely to be quite small (Bergin et al. 2012; Tubbs et al. 2020a), especially for any individual locus. These methods are often not powerful enough for locus discovery, so analysis of candidate SNPs or polygenic methods would be more appropriate at the current stage. Although the imputation of missing parents from half-siblings could increase power if included in our analyses (as show by Hwang et al., 2020), we chose not to include them in the current analyses to avoid any potential systematic differences in the parentally-provided environment between half-sibs and offspring from nuclear families.

Although we perform a genome-wide analysis in this paper as a demonstration of the method and for usage in future follow-up polygenic analyses, conditional models such as the one presented here are often not powerful enough for detection of loci with parental effects at the traditional GWAS significance level. For example, using the power calculations from Tubbs and colleagues (2020b), for a variant with an offspring and maternal genetic effect which both explain 0.1% of the variance in offspring phenotype, a sample size of less than 20,000 unrelated individuals would be necessary to detect a genome-wide significant effect with 80% power under a traditional GWAS model. However, under a conditional model, even under the best-case scenario when all individuals in each trio are genotyped, about 60,000 families would be necessary to achieve 80% power to detect a maternal effect and about 80,000 families would be needed to detect an offspring effect with the same power. Of course, power is also further reduced when some parents are missing and must be imputed. Depending on the goals of a particular study, the best approach when applying such conditional models with current sample size limitations may be to perform candidate SNP tests using known or suggestive loci identified by GWAS of unrelated individuals, which will decrease the multiple testing burden and increase power.

There are a number of potential limitations which serve to highlight future areas for extension and further application. First, our sample size is quite limited and may also suffer from selection bias such that family members who participated in the UKB could be significantly different across certain dimensions like personality or health than the remaining sample of unrelated individuals. Applying this procedure across available representative cohort studies would be ideal to reduce selection bias and increase sample size. Second, our imputation routine is applied SNP-by-SNP, which may not be ideal as it disregards information provided by variation around each SNP. Future work may explore the potential for haplotype-based imputation to increase imputation accuracy and power. Third, our model is simplified to assume the absence of assortative mating, dominance, or parent-of-origin effects, which may not reflect reality for most traits and some loci. Future work should more fully explore the consequences of violating these assumptions and potential solutions. Finally, while the linkage disequilibrium structure is preserved through imputation of missing parent genotypes, the correlation between SNPs is attenuated as a function of the allele frequencies. This makes the application of polygenic methods like univariate and bivariate LD score regression nontrivial. Future work will be necessary to determine the best method for addressing this attenuation to obtain unbiased estimates of variance explained and genetic correlations when utilizing summary statistics derived from observed and imputed genotypes.

Nevertheless, we have shown that our method of imputing missing parental genotypes can produce asymptotically unbiased estimates of parental genetic nurture effects and increase the effective sample size of datasets with missing parental genotypes. Ultimately, imputation of ungenotyped parents from their observed family members provides a powerful new tool for statistical geneticists interested in partitioning and estimating the effects of genetic nurture on human health and development.

## Acknowledgements

D.M.E. is funded by an Australian National Health and Medical Research Council Senior Research Fellowship (APP1137714) and NHMRC project grants (GNT1125200, GNT1157714, GNT1183074). This research has been conducted using the UK Biobank Resource under project ID number 28732.

