## Supplementary Material for "Modeling Parent-Specific Genetic Nurture in Families with Missing Parental Genotypes: Application to Birthweight and BMI"

**Data**

For *N* mother-offspring pairs, observed data include the offspring phenotype, *Y_o_*, and autosomal single nucleotide polymorphism (SNP) genotypes for both offspring, *X_o_*, and their mothers, *X_m_*. The coding of genotype assumes additivity (i.e. no dominance), and all genotypes and phenotypes are standardized to have mean 0 and variance 1.

As paternal genotypes, *X_f_*, are unavailable, they may be imputed for each SNP as

$$X_{f}^{'}=E(X_{f}|X_{o},X_{m})$$

Genotypic effects of *X_o_*, *X_m_*, and *X_f_’* may be tested for each SNP genotype or cumulatively through a polygenic risk score (PRS). Two potential methods for imputing the paternal PRS exist. First, one may impute the paternal genotype dosage from the offspring and maternal genotypes at each locus and apply standard PRS calculation methods to this derived dosage data using appropriate summary statistics. Otherwise, the maternal and offspring PRS may first be calculated and the offspring PRS regressed on the maternal PRS and the resulting residuals used as the imputed paternal PRS.

Of course, the same model may easily be adapted to the situation where the offspring and paternal genotypes are known, but the mother’s is unknown.

**Causal Model**

The offspring causal model is

$$Y_{o}=oX_{o}+mX_{m}+fX_{f}+\varepsilon_{o}$$

$$X_{o}=0.5\left( X_{m}+X_{f} \right)+\eta$$

where $\varepsilon_{o}$ is the error term for the phenotype and $\eta$ is the genotypic segregation variance such that

$$Var\left( \varepsilon_{o} \right)=\sigma^{2}$$

$$Var\left( \eta\right)=0.5$$

**Variances**

For the single SNP case, the variance of observed genotypes are equivalent to the expected heterozygosity, *H*, given the allele frequencies *p* and *q* at a given locus

$$Var\left( X_{o} \right)=Var\left( X_{m} \right)=Var\left( X_{f} \right)=2pq=H$$

Similarly, the variance of the imputed unobserved parental genotype at a given SNP is a function of *H*

$$Var\left( X_{f}^{'} \right)=Var\left( X_{m}^{'} \right)=\frac{2{H-H}^{2}}{4}$$

For ease of model non-centrality parameter (NCP) derivation, we standardize the genotypes relative to the observed genotypes, such that

$$Var\left( X_{o} \right)=Var\left( X_{m} \right)=Var\left( X_{f} \right)=1$$

$$Var\left( X_{f}^{'} \right)=Var\left( X_{m}^{'} \right)= \frac{2-H}{4}$$

The variance of the child phenotype is decomposed as

$$Var\left( Y_{o} \right)=\left( o^{2}+m^{2}+f^{2}+om+of \right)+\sigma^{2}=1$$

**Covariances**

We assume random mating, so that

$$Cov\left( X_{m},X_{f} \right)=0$$

The covariance between parent-offspring genotypes is

$$Cov\left( X_{o},X_{m} \right)=Cov\left( X_{o},X_{f} \right)=0.5$$

$$Cov\left( X_{o},X_{m}^{'} \right)=Cov\left( X_{o},X_{f}^{'} \right)=0.5$$

And the covariances between genotypes and phenotypes are

$$Cov\left( X_{o},Y_{o} \right)=o+0.5m+0.5f$$

$$Cov\left( X_{m},Y_{o} \right)=0.5o+m$$

$$Cov\left( X_{f},Y_{o} \right)=0.5o+f$$

$$Cov\left( X_{m}^{'},Y_{o} \right)=0.5o+mVar(X_{m}^{'})$$

$$Cov\left( X_{f}^{'},Y_{o} \right)=0.5o+fVar(X_{f}^{'})$$

The covariance between the imputed parental genotype and the corresponding observed genotype is equal to its variance

$$Cov\left( X_{f}^{'},X_{f} \right)=Cov\left( X_{m}^{'},X_{m} \right)=Var\left( X_{f}^{'} \right)=Var\left( X_{m}^{'} \right)$$

**Power calculation under linear regression analysis**

When offspring, maternal, and paternal genotypes are all observed, the linear model is

$$Y_{o}=oX_{o}+mX_{m}+fX_{f}+\varepsilon_{o}$$

The fixed effects *o, m,* and *f* are estimated by ordinary least squares (OLS) regression, where the vector of asymptotic parameter estimates, $\beta$, is given by

$$\hat{\beta}={\Sigma_{XX}}^{-1}\Sigma_{XY}$$

where ${\Sigma_{XX}}^{-1}$ is the inverse of the variance-covariance matrix of independent variables included in the model and $\Sigma_{XY}$ is a column vector of covariances between the dependent variable and each independent variable.

Similarly, the residual variance, $\sigma^{2}$ for a given model is:

$$\sigma^{2}=1-{{\Sigma_{YX}\Sigma}_{XX}}^{-1}\Sigma_{XY}$$

where $\Sigma_{YX}$ is row vector of covariances between *y* and the independent variables.

Given a sample size of *n* sibling pairs, the covariance matrix of beta estimates $\Sigma_{B}$ is

$$\Sigma_{B}=\sigma^{2}\frac{{\Sigma_{XX}}^{-1}}{n}$$

where the square root of the diagonal elements in this matrix give the standard errors (SE) for the corresponding beta estimates.

Therefore, a non-centrality parameter (NCP) may be calculated for each beta estimate, $\hat{\beta}$, as

$${NCP}_{\hat{\beta}}=\left( \frac{\hat{\beta}}{{SE}_{\hat{\beta}}} \right)^{2}$$

**Observed Genotypes Model**

We first consider the model where all genotypes are observed, so that independent variables include *X_o_*, *X_m_*, and *X_f_* . Solving for $\hat{\beta}$ gives unbiased estimators such that

$$\hat{o}=o$$

$$\hat{m}=m$$

$$\hat{f}=f$$

and an expected error variance of:

$$\sigma^{2}=1-\left[ o^{2}+m^{2}+f^{2}+om+of \right]$$

with the NCP for each beta estimate being:

$${NCP}_{\hat{o}}=\frac{\hat{o}^{2}n}{2\sigma^{2}}$$

$${NCP}_{\hat{m}}=\frac{\hat{m}^{2}2n}{3\sigma^{2}}$$

$$\mathrm{NCP}_{\hat{f}}=\frac{\hat{f}^{2}2n}{3\sigma^{2}}$$

**Single Missing Parental Genotype Model**

When the genotypes of an offspring and one parent are observed, the genotype of the other parent may be imputed. For the derivation, we consider the case when the mother and offspring genotypes are known and the father’s is imputed, thus *X_o_*, *X_m_*, and *X_f_’* are included as independent variables. Solving for the vector of regression coefficients provides unbiased estimates of the genetic effects

$$\hat{o}=o$$

$$\hat{m}=m$$

$$\hat{f}=f$$

and an expected error variance of

$$\sigma^{2}=1-\left[ o^{2}+m^{2}+f^{2}Var(X_{f}^{'})+om+of \right]$$

with the NCP for each beta estimate being

$${NCP}_{\hat{o}}=\frac{\hat{o}^{2}n(3Var\left( X_{f}^{'} \right)-1)}{\sigma^{2}4Var(X_{f}^{'})}$$

$${NCP}_{\hat{m}}=\frac{\hat{m}^{2}n(3Var\left( X_{f}^{'} \right)-1)}{\sigma^{2}(4Var\left( X_{f}^{'} \right)-1)}$$

$$\mathrm{NCP}_{\hat{f}}=\frac{\hat{f}^{2}n(3Var\left( X_{f}^{'} \right)-1)}{3\sigma^{2}}$$

**Method for Combining Effect Estimates from Sibship-based and Parent-Offspring Duo-based Missing Parent Imputation Models**

Our approach is to perform two separate mixed model analyses: (1) families with 0 or 1 missing parent, (2) families with 2 missing parents. We can obtain estimates for both paternal and maternal effects from (1), together with a covariance matrix for the estimates. For (2), the 2 parents have the same imputed genotypes, and so we enter only one of them and interpret the estimate for the effect of that imputed genotype to be an estimate of the sum of the paternal and maternal effects. We can then combine the estimates from (1) and (2) into a single set of estimates for the two parental effects.

From (1), we obtain separate estimates of paternal and maternal effects: *f* and *m,* respectively. We also obtain their covariance matrix. Let the estimation variance of *f* be *V_f_*, the estimation variance of *m* be *V_m_*, and the estimation covariance between *f* and *m* be *C_fm_*. From these outputs we estimate the sum of the effects of the two parents as *s = f + m*, with estimation variance *V_f_ +V_m_+2C_fm_*. Similarly, we estimate the difference of the effects of the two parents as *d = f – m*, with estimation variance *V_f_ +V_m_ – 2C_fm_*. The estimation covariance between *s* and *d* is *V_f_ –V_m_*.

From (2), we only obtain one estimate, which represents an estimate of the combined parental effects, which we call *c,* with estimation variance *V_c_*.

Since c and s are both estimates of the total effects of the two parents, we can simply combine them into a single estimate *t* by inverse variance weighting such that

$$t=\frac{\left( \frac{s}{\left( V_{f}+V_{m}+{2C}_{fm} \right)} \right)+\left( \frac{c}{V_{c}} \right)}{\left( \frac{1}{\left( V_{f}+V_{m}+{2C}_{fm} \right)}+\frac{1}{V_{c}} \right)}$$

$$=\frac{sV_{c}+c\left( V_{f}+V_{m}+2C_{fm} \right)}{\left( V_{f}+V_{m}+{2C}_{fm}+V_{c} \right)}$$

with variance

$$V_{t}=\frac{1}{\left( \frac{1}{\left( V_{f}+V_{m}+{2C}_{fm} \right)}+\frac{1}{V_{c}} \right)}$$

$$=\frac{\left( V_{f}+V_{m}+{2C}_{fm} \right)V_{c}}{V_{f}+V_{m}+{2C}_{fm}+V_{c}}$$

If we have two parameters *a* and *b* (representing the true paternal and maternal effects), and we have an estimate of their sum *t* *= (a+b)* and an estimate their difference *d = (a–b)*, then it would seem reasonable to estimate *a* by *(t+d)/2*, and *b* by *(t–d)/2*.

The variance of *(t+d)/2* should be

$$Var\left( \frac{t+d}{2} \right)=\frac{Var\left( t+d \right)}{4}$$

$$=\frac{Var\left( t \right)+Var\left( d \right)+2Cov\left( t,d \right)}{4}$$

$$=\frac{{{\left( V_{f}+V_{m}+{2C}_{fm} \right)V_{c}+(V}_{f}+V_{m}-{2C}_{fm})(V}_{f}+V_{m}+{2C}_{fm}+V_{c})+2V_{c}(V_{f}-V_{m})}{4(V_{f}+V_{m}+{2C}_{fm}+V_{c})}$$

$$=\frac{\frac{V_{f}+V_{m}-{2C}_{fm}+V_{c}(3V_{f}-V_{m}+{2C}_{fm})}{V_{f}+V_{m}+{2C}_{fm}+V_{c}}}{4}$$

Similarly, the variance of *(t–d)/2*should be

$$Var\left( \frac{t-d}{2} \right)=\frac{Var\left( t-d \right)}{4}$$

$$=\frac{Var\left( t \right)+Var\left( d \right)-2Cov\left( t,d \right)}{4}$$

$$=\frac{{{\left( V_{f}+V_{m}+{2C}_{fm} \right)V_{c}+(V}_{f}+V_{m}-{2C}_{fm})(V}_{f}+V_{m}+{2C}_{fm}+V_{c})-2V_{c}(V_{f}-V_{m})}{4(V_{f}+V_{m}+{2C}_{fm}+V_{c})}$$

$$=\frac{\frac{V_{f}+V_{m}-{2C}_{fm}+V_{c}(3V_{m}-V_{f}+{2C}_{fm})}{V_{f}+V_{m}+{2C}_{fm}+V_{c}}}{4}$$

The above derivation requires *Cov(t,d)*, which can be shown to be

$$Cov\left( t,d \right)=Cov\left( \frac{V_{c}\left( f+m \right)}{V_{f}+V_{m}+{2C}_{fm}+V_{c}},(f-m) \right)$$

$$=\frac{V_{c}(V_{f}-V_{m})}{V_{f}+V_{m}+{2C}_{fm}+V_{c}}$$

Finally, the covariance between the estimates of *a* and *b* is

$$\frac{Cov\left( \left( t+d \right),(t-d) \right)}{4}=\frac{V_{t}-V_{d}}{4}$$

$$=\frac{V_{c}(V_{f}+V_{m}+{2C}_{fm})}{4\left( V_{f}+V_{m}+{2C}_{fm}+V_{c} \right)}-\frac{V_{f}+V_{m}-{2C}_{fm}}{4}$$

Clearly, we would wish to perform separate test for the effects of the two parents, *a* and *b*. However, it may also make sense to perform tests of *t* and *d*, if the hypothesis of different paternal and maternal effects is reasonable. Thus, we could first test for *d*. If this is significant, then we proceed to test for *a* and *b* separately. If this is not significant, we can proceed to test for *t*.
