## Supplementary Figures for "Modeling Parent-Specific Genetic Nurture in Families with Missing Parental Genotypes: Application to Birthweight and BMI"

**Supplementary Figure 1. Power for Imputed vs Observed Model**

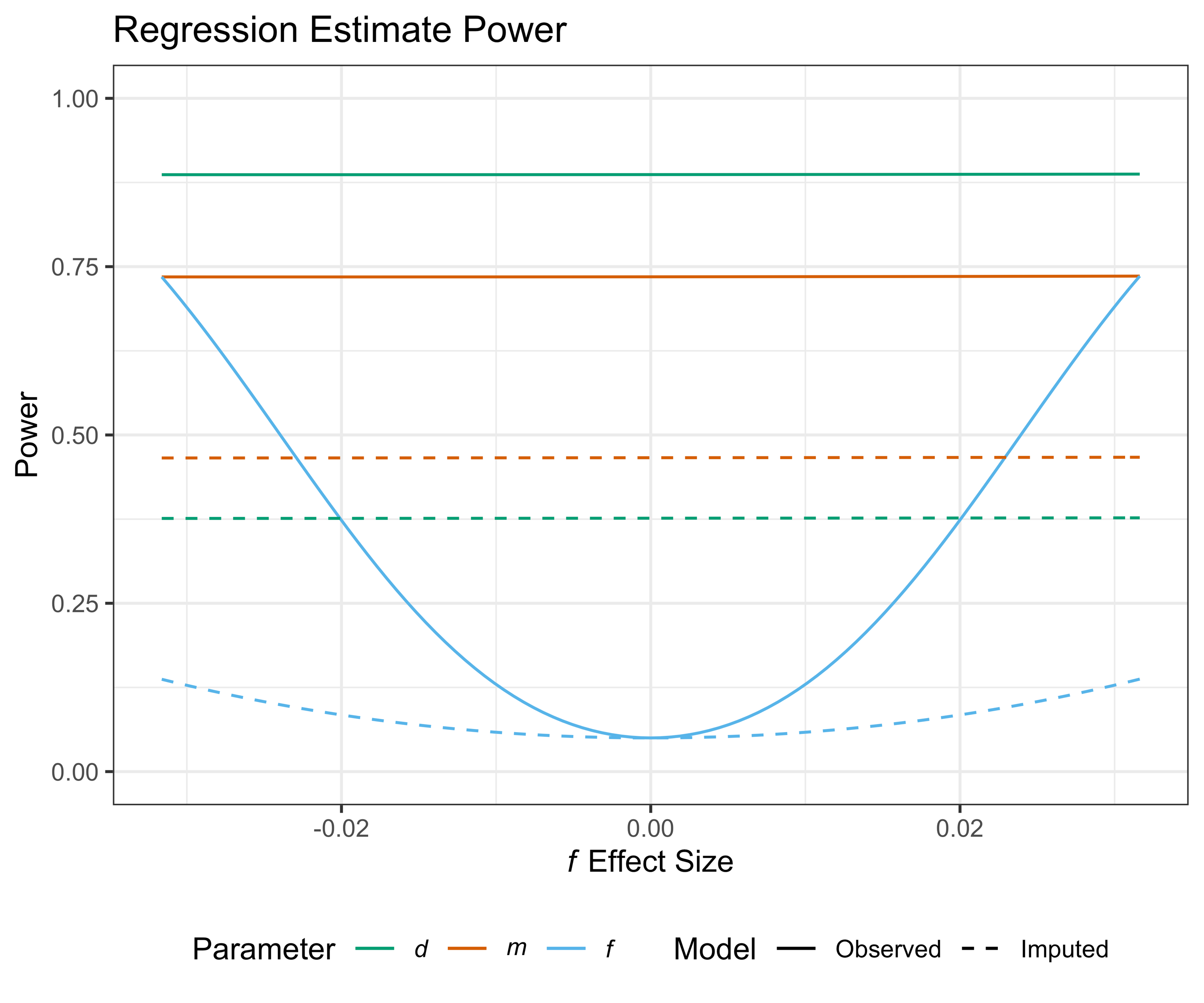

*Note: Power for each parameter is defined by the area above the NCP (as derived in the supplementary material) under the non-central chi square distribution with one degree of freedom and an alpha significance level of 0.05. The sample size was constrained to 10,000 family units. The offspring effect, d, and the maternal effect, m, were set to explain 0.2% and 0.1% of the variance in the offspring phenotype, respectively. The paternal effect, f, explained between 0 and 0.1% of the variance, with effect sizes in the positive and negative direction. The Observed Model is that in which the offspring phenotype is regressed on observed offspring, maternal, and paternal genotypes. The Imputed Model differs only in that the paternal genotype imputed from mother-child duos is included rather than the observed paternal genotype. Thus, when one parameter is being tested, the other two are also included in the model. Since the father’s genotype is included in the model estimation procedure and its estimate is asymptotically unbiased, the power to detect an effect of a maternal or offspring effect is independent of the true paternal effect size, but rather depends on the imputation accuracy, which is a function of the population allele frequency.*

**Supplementary Figure 2. Manhattan Plots of Offspring and Maternal Genetic Effects**

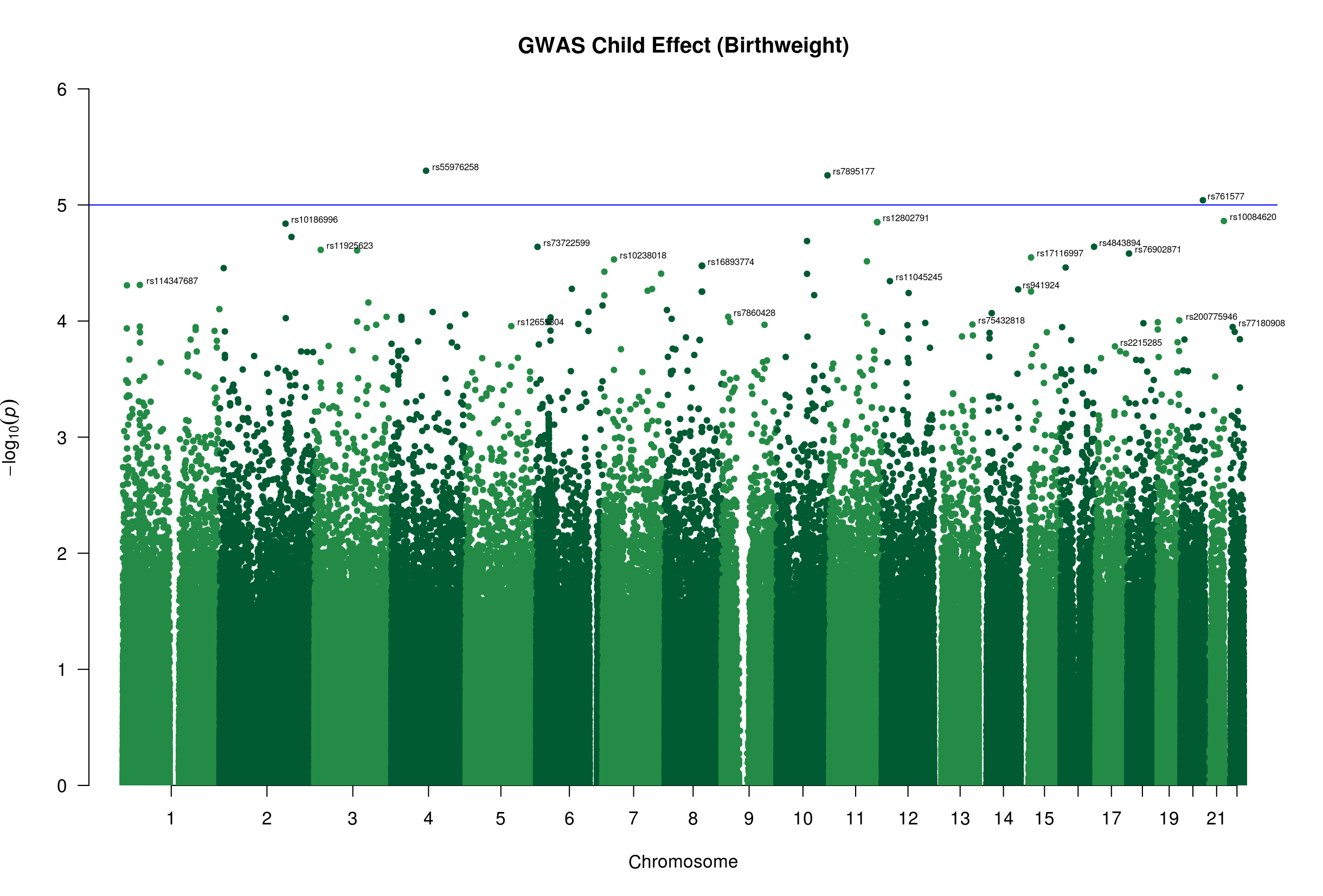

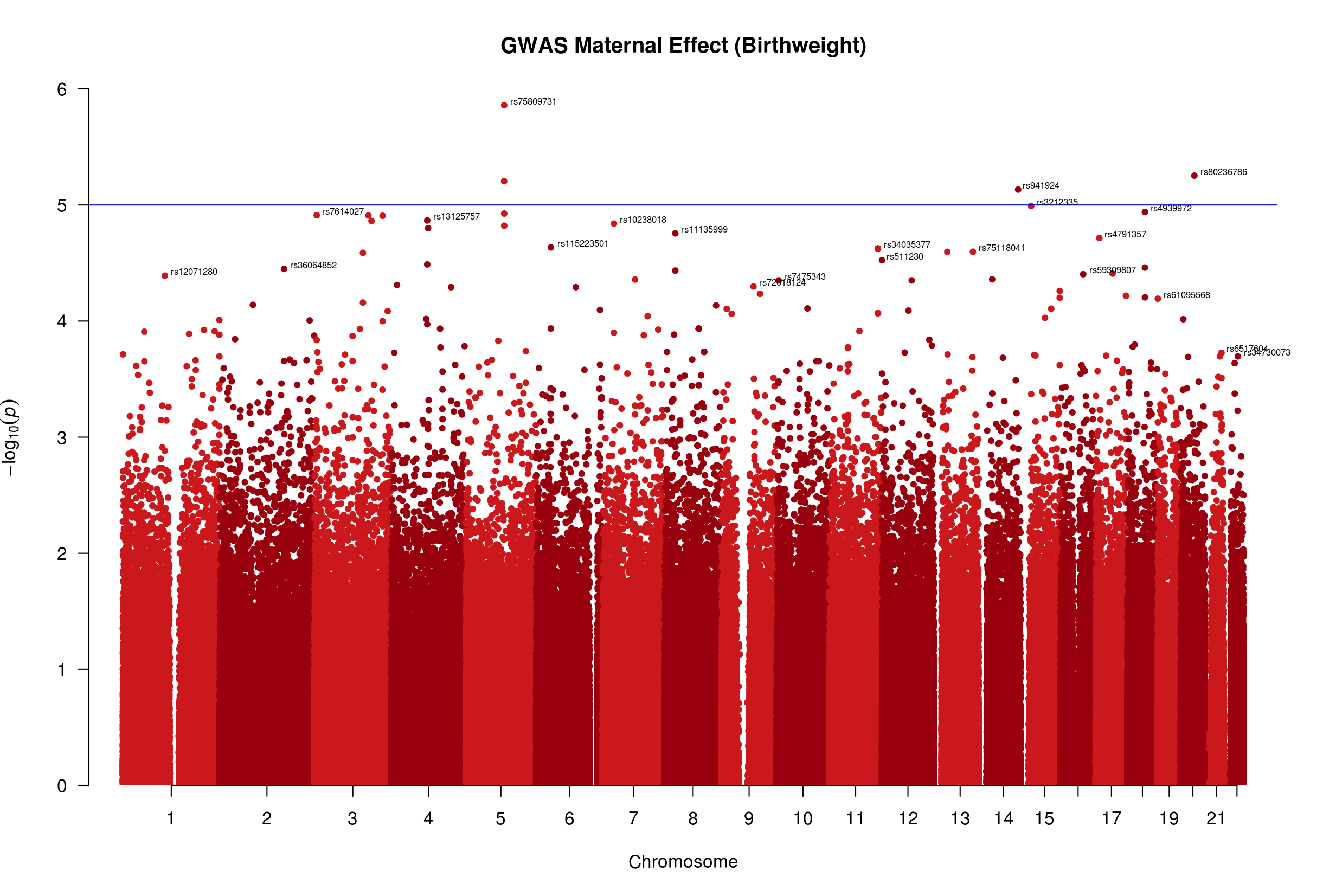

*Manhattan plots for offspring and maternal genetic effects on offspring birthweight. Blue horizontal line represents a suggestive genome-wide significance level of 1e-5. P-values are corrected through genomic control with genomic inflation factor* $\lambda=1.04$ *for offspring effect and* $\lambda=1.02$ *for maternal effect. The SNP with the highest p-value for each chromosome is labeled with its RSID.*

**Supplementary Figure 3. QQ Plots for Birthweight and BMI GWAS**

a. Birthweight

Child Effect	Maternal Effect

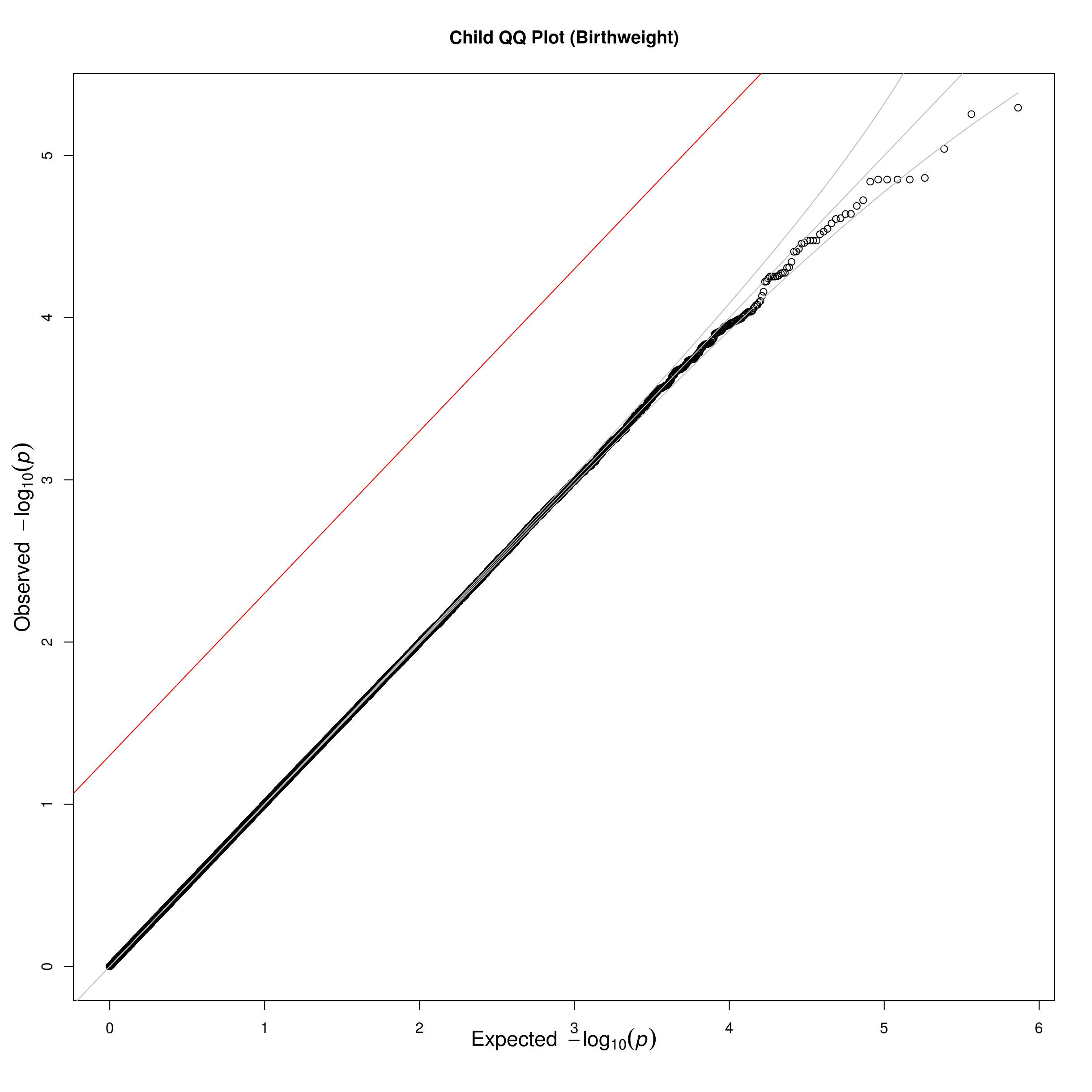

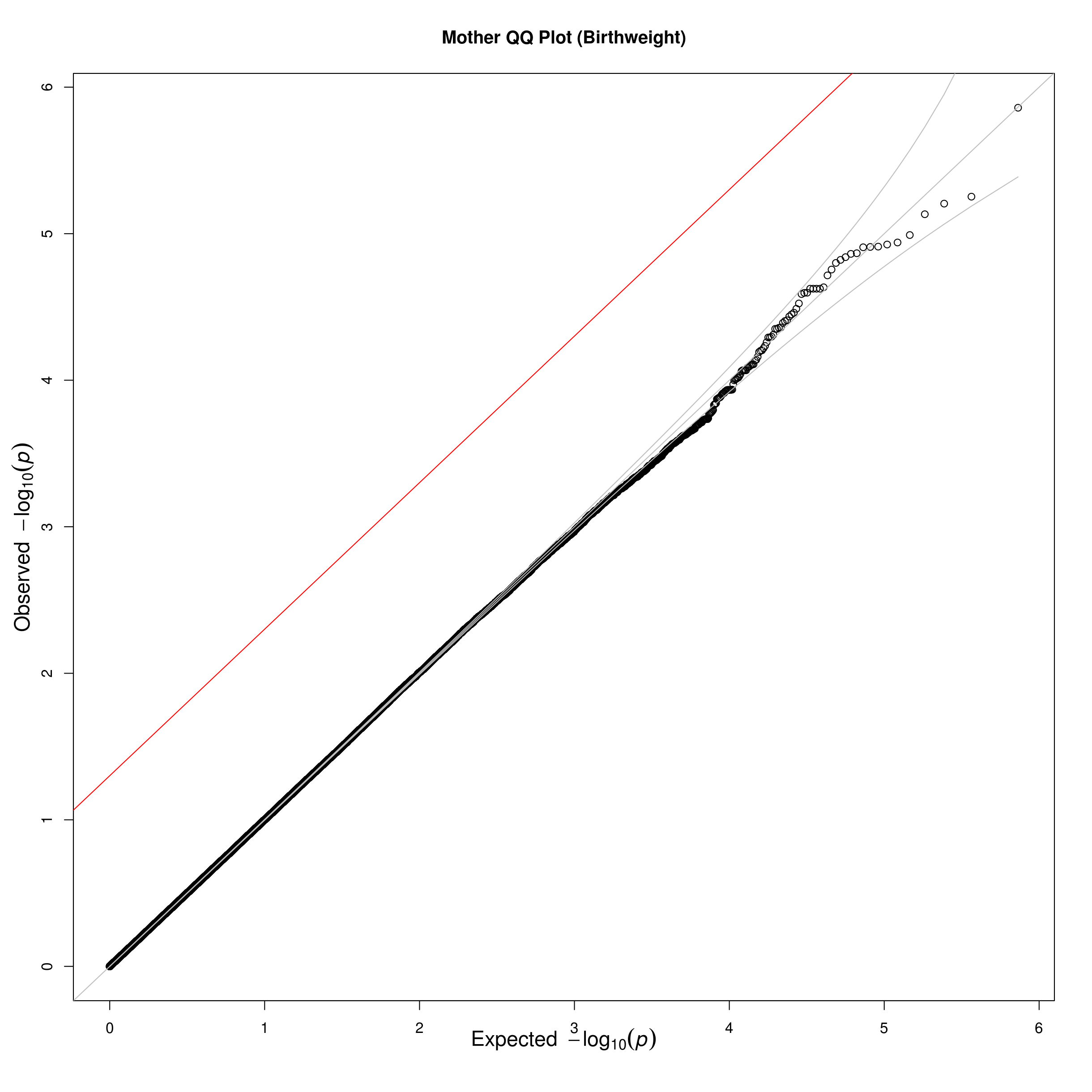

b. BMI

Child Effect	Maternal Effect	Paternal Effect

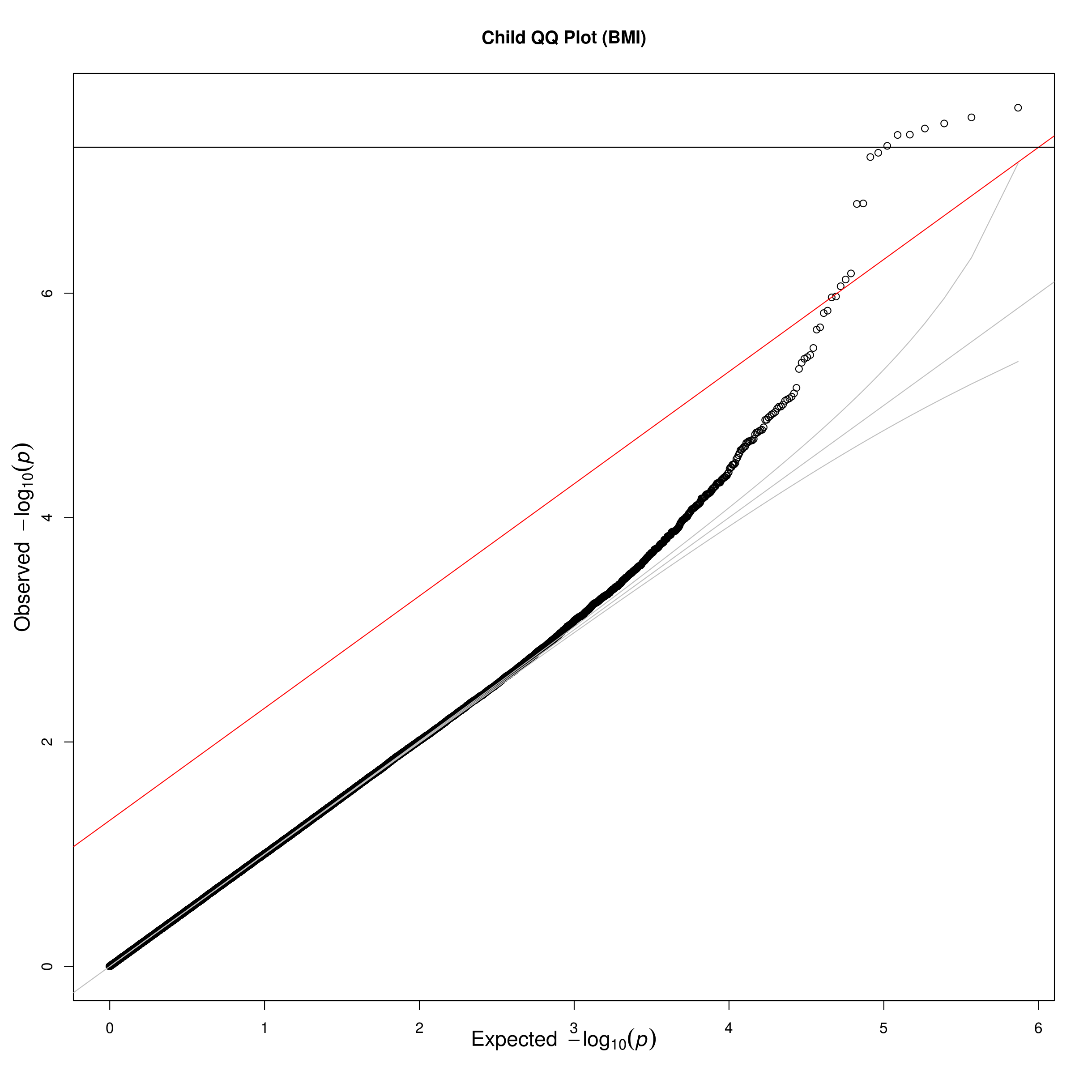

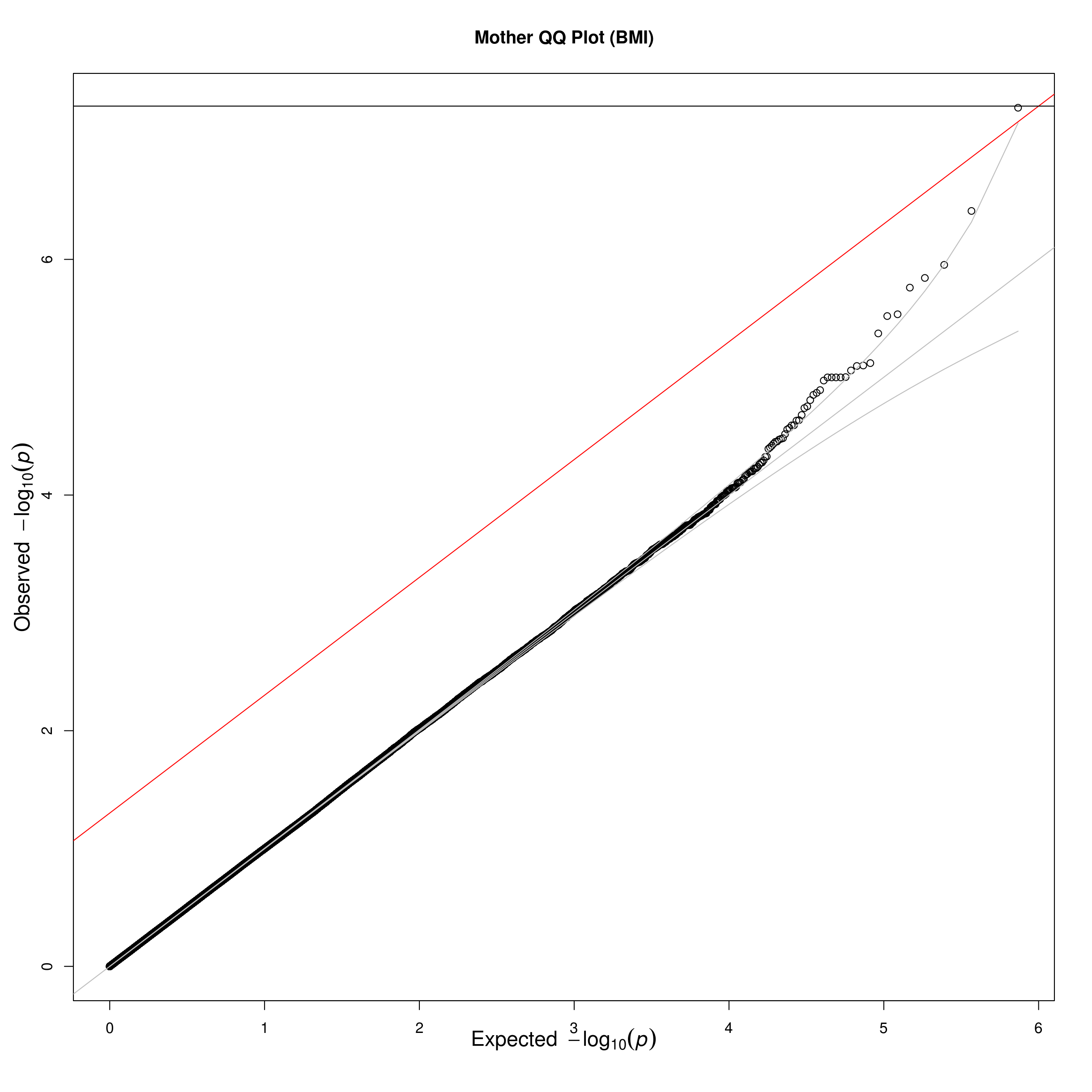

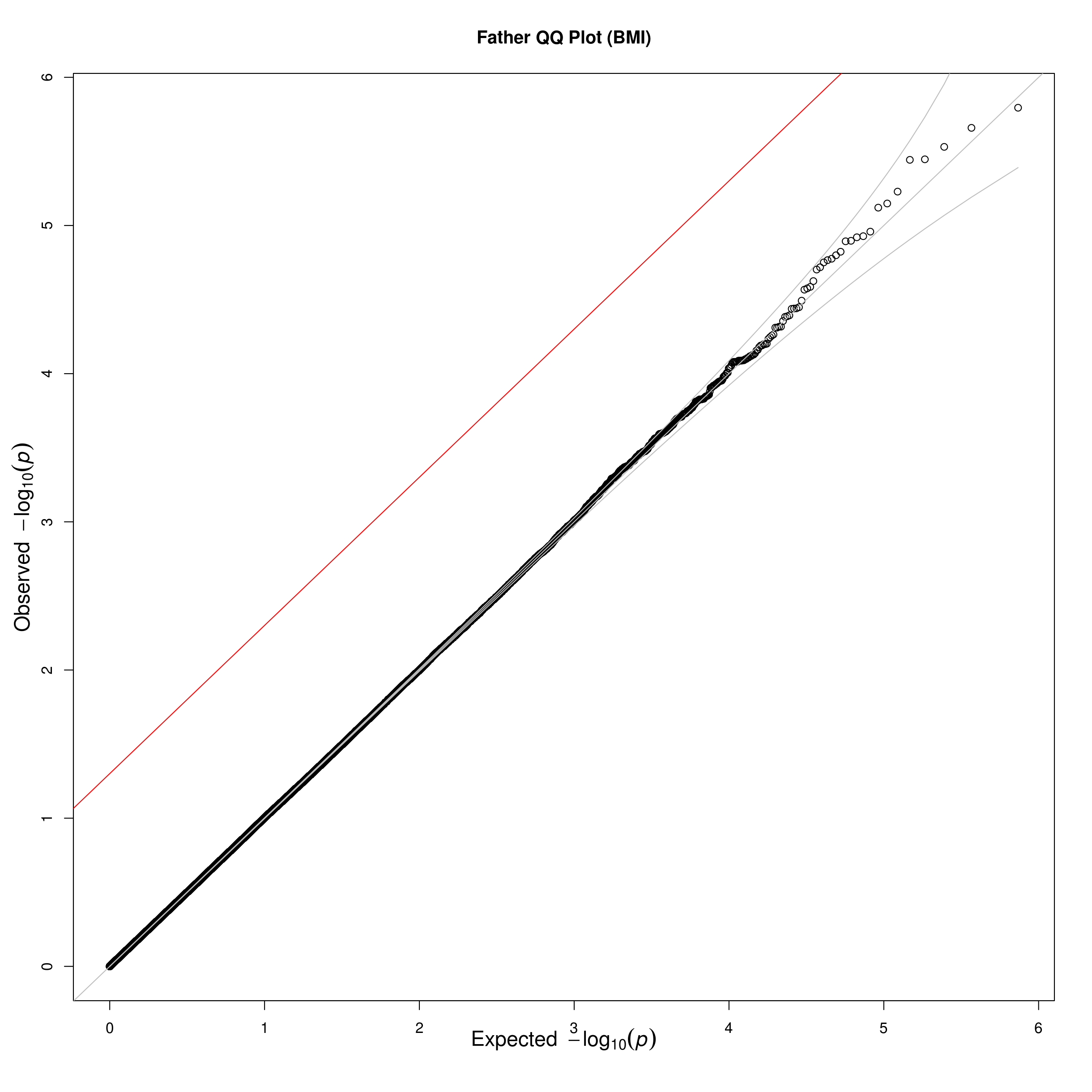

*QQ plots for parental and child genetic effects on birthweight and BMI. Diagonal red line represents the false discovery rate threshold with an alpha level of 0.05. The horizontal black line represents the Bonferroni corrected threshold of 5e-8. P-values are corrected through genomic control with genomic inflation factor* $\lambda=1.04$ *for offspring effect on birthweight and* $\lambda=1.02$ *for maternal effect on birthweight. For BMI,* $\lambda=1.04$ *for offspring effect,* $\lambda=1.04$ *for maternal effect, and* $\lambda=1.03$ *for paternal effect.*

**Supplementary Figure 4. Manhattan Plots for Offspring, Maternal, and Paternal Genetic Effects on BMI**

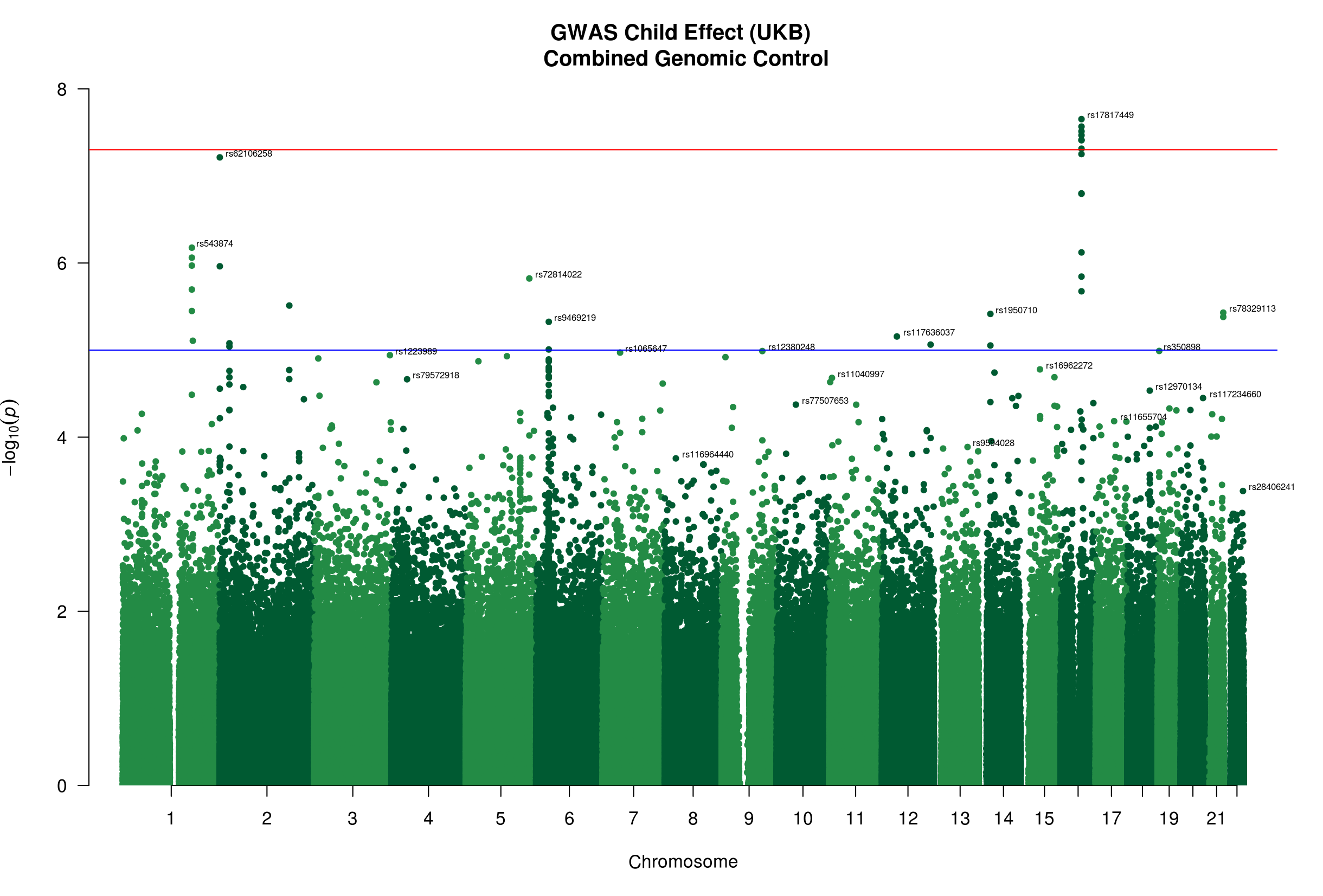

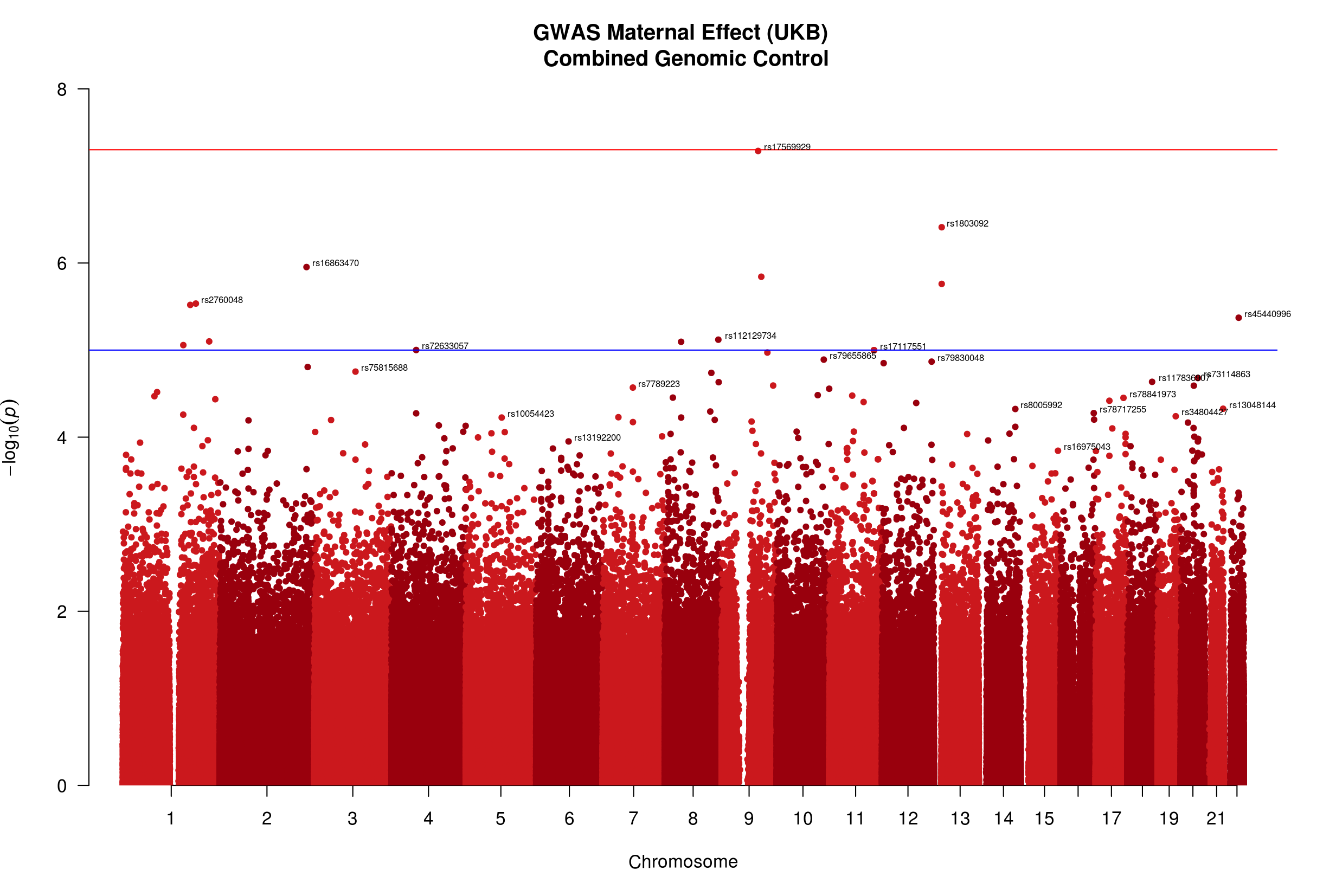

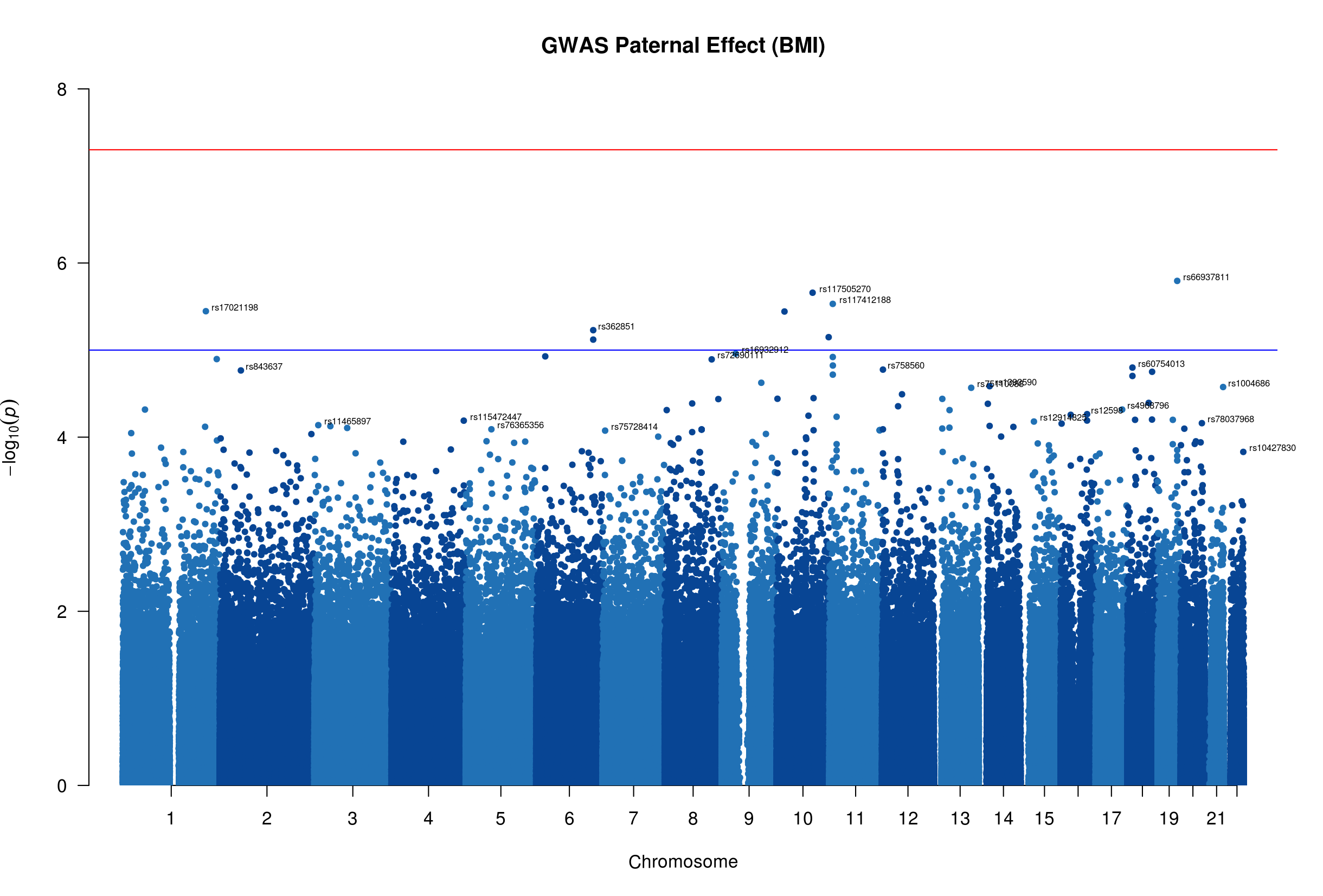

*These manhattan plots show the -log p-values (after applying genomic control correction with* $\lambda=1.04$ *for offspring effect,* $\lambda=1.04$ *for maternal effect, and* $\lambda=1.03$ *for paternal effect) for observed SNPs in our imputation model estimating child, maternal, and paternal genetic effects on offspring birthweight in the UK Biobank. The red horizontal line indicates the traditional GWAS p-value threshold of 5e-8 and the blue horizontal line indicates a suggestive significance threshold of 1e-5. The SNP with the highest p-value for each chromosome is labeled with its RSID.*
